# Cytomegalovirus infection in newborn mice alters cerebellar development by lengthening G1/S phases of cerebellar granule cell precursors during postnatal cerebellar development

**DOI:** 10.1101/2022.10.02.510512

**Authors:** Cathy Yea Won Sung, Mao Li, Stipan Jonjic, Veronica Sanchez, William J Britt

## Abstract

Human cytomegalovirus (HCMV) infection of the developing central nervous system (CNS) in infants infected *in utero* can lead to a variety of neurodevelopmental disorders. Although the link between HCMV infection and neurodevelopmental deficits is widely recognized, underlying mechanisms leading to altered neurodevelopment remain poorly understood. We have previously described a murine model of congenital HCMV infection in which murine CMV (MCMV) spreads hematogenously and establishes a focal infection in the brain of newborn mice. Infection results in the disruption of cerebellar cortical development characterized by reduced cerebellar size, but paradoxically, an increase in the number of cerebellar granule cell precursors (GCPs) in the external granular layer (EGL) of the cerebellar cortex. This increased number of GCPs in the EGL is associated with abnormal cell cycle progression and decreased GCP migration from EGL and IGL. In the current study, we demonstrated that MCMV infection led to prolonged G1- and S-phases of the GCP cell cycle and increased cell cycle exit. Treatment with TNFα neutralizing antibody partially normalized the cell cycle progression of GCPs. Collectively, our results argue that inflammation can alter GCP proliferation and lead to premature exit from the cell cycle resulting in reduced cerebellar size in MCMV-infected mice. These findings provide insight into mechanisms of altered brain development of fetuses infected with HCMV and possibly, other infectious agents that induce inflammation during neurodevelopment.

## Introduction

Congenital human cytomegalovirus (HCMV) infection is a major cause of morbidity in infants and children throughout the world, affecting 0.2-1.2% of all live births in the United States. Among infected infants, approximately 5-15% of newborn infants (~3000/yr in US) will develop long-term neurodevelopmental sequelae. Studies in fetuses with congenital HCMV infection using magnetic resonance fetal imaging have documented altered brain morphogenesis including ventriculomegaly, microcephaly, lissencephaly, cortical dysplasia, periventricular calcification, and cerebellar hypoplasia (Malinger et al., 2003). Histopathological studies in the brains of infected fetuses and necropsies of newborn infants with congenital HCMV infection have revealed increased monocytic infiltration, reactive gliosis, and foci of CD8+ T cell aggregates (Gabrielli et al., 2012). Tissue damage was frequently observed not only in regions with HCMV-infected cells but also in regions without evidence of virus infection suggesting that virus-induced inflammation could also contribute to the damage to the CNS (Gabrielli et al., 2012). While histopathological changes associated with congenital HCMV infection are well-described, the pathogenesis of HCMV-induced damage in the CNS remains undefined. We developed a mouse model in which newborn mice are infected intraperitoneally with a non-lethal dose of murine CMV (MCMV), a virus with similar genetic background and replication program to HCMV. Following infection, MCMV infects peripheral organs and subsequently spreads hematogenously to the CNS. Although this model does not recapitulate *in utero* transmission of the virus, it takes advantage of results from previous studies that have shown that the newborn mouse is neurodevelopmentally similar to a late 2nd trimester human fetus (Bortolussi et al., 2014; Moulden et al., 2021).

The cerebellum in rodents develops postnatally reaching its maturity at approximately postnatal day (PNd) 21 (Goldowitz and Hamre, 1998; Millen et al., 1994). Cerebellar granule cell precursors (GCPs) are the most abundant cell type in the developing cerebellum and are responsible for normal cellular positioning of the laminar structure of the cerebellar cortex that is composed of the external granule layer (EGL), molecular layer (ML), Purkinje cell layer (PCL), and internal granule layer (IGL). GCPs are mitotically active in the outer EGL (oEGL) where proliferation is maximal at PNd8 in response to sonic hedgehog (SHH) produced by Purkinje cells (PCs) (Behesti and Marino, 2009). After multiple rounds of proliferation, GCPs exit the cell cycle, move into the deeper layer of the EGL (pre-migratory layer, inner EGL, iEGL), and differentiate into mature granule cells (GCs). GCs then migrate radially along the Bergmann glial axons in the ML, passing the PCL containing the soma of PCs and Bergmann glia, and form the IGL, which is the final position of GCs (Goldowitz and Hamre, 1998).

We have previously demonstrated that the morphogenesis of the cerebellum is altered in MCMV-infected mice (Koontz et al., 2008; Kosmac et al., 2013). Abnormalities in cerebellar development during MCMV infection include globally altered morphogenesis (e.g. decreased cerebellar size, area, weight, and foliation) and changes in cortical structures (e.g. thicker EGL, decreased thickness of ML and IGL). These changes in cerebellar development were not linked directly to virus-induced cytopathology as the CNS infection in this model is focal with no histologic evidence of necrosis and/or apoptosis of resident cells in the cerebellum (Koontz et al., 2008). Cerebellar hypoplasia in MCMV-infected animals was associated with reduced GCP proliferation in the EGL and delayed migration from the EGL to the IGL (Koontz et al., 2008). During this developmental period, robust inflammatory response, such as cytokine release (e.g., interferon-stimulated genes and TNFα), reactive gliosis, recruitment of inflammatory monocytes/macrophages, neutrophils, natural killer (NK) cells, and CD8+ T cells can be readily detected in the brains of MCMV-infected mice, suggesting that the host immune response induced by virus infection contributed to the global and symmetric changes in the cerebellum as well as in the hearing organ, the cochlea (Bantug et al., 2008; Cheeran et al., 2001; Koontz et al., 2008; Kosugi et al., 2000; Sung et al., 2019; van den Pol et al., 2007). Consistent with a mechanism of virus-induced immunopathology, we have demonstrated that corticosteroid or anti-TNFα neutralizing antibody treatment partially corrected the abnormalities in the developing CNS, while minimally impacting the level of virus replication in the CNS (Kosmac et al., 2013; Seleme et al., 2017; Sung et al., 2019). In contrast to these previous studies, a more recent study utilizing intracerebral inoculation of embryonic mice demonstrated damage to the developing CNS that appear to be directly attributable to virus replication and cytopathology at early stages of neurodevelopment (Zhou et al., 2022).

A variety of mechanisms can lead to impaired neurogenesis and the development of microcephaly or cerebellar hypoplasia (Cremisi et al., 2003). Pediatric patients or fetuses with Down syndrome (DS) exhibit brain hypoplasia or microcephaly, similar to neurologic manifestations of congenital HCMV infection. Studies have shown reduced neurogenesis in the EGL of the cerebellum, hippocampus, and ventricular zone (VZ) in fetuses with DS and in engineered murine models of DS (Contestabile et al., 2007; Guidi et al., 2011). Specifically, in Ts65Dn mice, GCP cell cycle was significantly delayed due to prolonged G1- and G2-phases and impaired neurogenesis attributed to decreased responsiveness to the mitogenic factor SHH during postnatal cerebellar development (Contestabile et al., 2009; Roper et al., 2006).

A number of studies in the developing cerebral cortex have demonstrated that neural progenitor cell (NPC) proliferation and brain size are largely influenced by cell cycle length, mainly due to the correlation between G1 lengthening and NPC differentiation status (Calegari et al., 2005; Cremisi et al., 2003; Dehay and Kennedy, 2007; Farkas and Huttner, 2008; Lange and Calegari, 2010). Growing evidence suggests that decreasing the length of G1 can lead to an inhibition of neurogenesis and expansion of progenitor pool, while lengthening of the G1-phase leads NPCs to transition from proliferative symmetric division to asymmetric neurogenic division (Artegiani et al., 2011; Kaldis and Richardson, 2012; Lange et al., 2009; Mitsuhashi et al., 2001; Pilaz et al., 2009). These studies support the hypothesis that the length of G1 is a critical determinant of cell differentiation.

The developing cerebellum exhibits unique features that are distinct from the cerebral cortex. In rodents, GCPs are generated from the rhombic lip (RL) between E12.5 and E15.5, migrate to the cerebellar anlage forming the EGL, and continue to actively proliferate throughout the postnatal development until about 21 days after birth, unlike NPCs in the developing cerebral cortex in which proliferation and differentiation occurs embryonically (Gao and Hatten, 1993; Greig et al., 2013). In addition, GCPs in the developing cerebellum predominantly undergo symmetric division, switching from non-terminal symmetric division to terminal symmetric division as development proceeds, regulated primarily by SHH that is secreted by PCs (Consalez et al., 2020; Espinosa and Luo, 2008; Nakashima et al., 2015; Yang et al., 2015). However, the correlation between different stages of postnatal cerebellar development and the cell cycle parameters of GCPs have not been characterized extensively in control and diseased brains, such as in MCMV-infected mice. Furthermore, many studies that have described the impact of insults such as inflammation on cerebellar development have been carried out *in vitro* in tissue explants and/or isolated resident cells of the cerebellum and not in *in vivo* models.

In the current study, we utilized well described methodologies to define cell cycle parameters to investigate the reduced proliferation of GCPs in the developing cerebellum of control and MCMV-infected mice. Our results demonstrated that MCMV infection induced robust inflammation in the developing cerebella of newborn mice and lengthened the duration of GCP cell cycle by prolonging G1- and S-phases compromising GCP proliferation. In addition, we observed that GCPs prematurely exited cell cycle resulting in decreased cellularity of the IGL and cerebellar hypoplasia, a characteristic of cerebella from infected mice that we have previously reported (Koontz et al., 2008). Decreasing inflammation in this model by treatment with anti-TNFα neutralizing antibody partially corrected the cell cycle abnormalities in GCPs of MCMV-infected mice and, as we have previously shown, normalized some of the morphologic abnormalities of the cerebella of mice infected with MCMV (Seleme et al., 2017). Our findings in this study provide additional evidence of the impact of virus-induced inflammation on neurodevelopment.

## Materials and Methods

### Ethics statement

All animal protocols (APN9351) were approved from Institutional Animal Care and Use Committee (IACUC) of the University of Alabama at Birmingham (UAB). Mice were euthanized by carbon dioxide (CO2) asphyxiation followed by cervical dislocation for adult mice and decapitation for mice younger than PNd12. All experimental procedures were approved from IACUC and the UAB Animal Resource Program (ARP) and are in compliance with guidelines for care and the use of laboratory animals to harvest tissues for this project.

### Animals and MCMV infection

Pathogen-free BALB/c mice were purchased from Jackson Laboratories (Bar Harbor, ME) and housed under specific pathogen-free conditions. MCMV virus (Smith strain repaired M128) stocks were propagated in M2-10B4, mouse bone marrow stromal cell line (ATCC, CRL-1972), and harvested at the peak of cytopathic effect (Lutarewych, 1997; Jordan, 2011). Aliquots of virus stocks were stored in −80°C. Infectious virus titers were measured by standard plaque assay in mouse embryonic fibroblast (MEF). Both male and female newborn mouse pups were infected with 500 plaque forming unit (PFU) of MCMV virus diluted in sterile phosphate-buffered saline (PBS). Sex has not been shown to be a determinant in MCMV-induced CNS disease or CNS disease associated with HCMV infection of the developing CNS of humans. Virus injection was performed intraperitoneally (i.p.) within 12 hrs following birth as previously described (Koontz et al., 2008).

### TNFα neutralizing antibody (TNF-NAb) treatment

On postnatal day (PNd) 3-7, pups were treated daily by i.p. injection with *InVivoMAb* rat IgG1 Isotype control anti-trinitrophenol (TNP) (BioXCell, Lebanon, NH) or *InVivoMAb* anti-mouse TNFα neutralizing antibody (Rat IgG1, clone. XT3.11, BioXCell, Lebanon, NH) at 500 μg/mouse/day diluted in 1X PBS. MCMV-infected and uninfected, control mice that were not treated with either TNF-NAb or isotype control antibody but received a vehicle injection of sterile 1X PBS. On PNd8, mouse pups were sacrificed and following exhausative perfusion carried out by insertion of a needle into the left ventricle followed by perfusion with ice-cold 1X PBS, organs including brains were harvested and prepared for appropriate downstream assays.

### Quantitation of virus genome copy number and gene expression

Mice were sacrificed at various time points, perfused as described above, and organs including brains or dissected cerebellum were harvested. DNA and RNA were isolated from the cerebellum using E.Z.N.A Total RNA kit (Omega Bio-tek, Norcross, GA) with minor modifications in the manufacturer’s protocol. Quantitative PCR (qPCR) was performed to detect amplification of viral immediate early-1 (IE-1) gene exon 4 from total DNA using forward primer (5’-GGC TCC ATG ATC CAC CCT GTT A-3’), reverse primer (5’-GCC TTC ATC TGC TGC CAT ACT-3’) and probe (5’-AGC CTT TCC TGG ATG CCA GGT CTC A-3’) labeled with FAM and TAMRA. A standard curve was generated from serial dilutions of IE-1 exon 4 cloned into pcDNA3. qPCR was performed using TaqMan Gene Expression master mix (ThermoFisher Scientific, Wiltham, MA). Cerebellar samples were run in duplicates on the StepOne Plus Real-Time PCR system (Applied Biosystems, Life Technologies, Foster City, CA). Viral genome copy numbers were expressed as log(10) genome copies per milligram (mg) of tissue. For quantitative reverse transcription PCR (RT-PCR) assays, the Invitrogen Superscript III First strand synthesis kit (Thermo Fisher Scientific, Waltham, MA) was used to synthesize cDNA from total RNA according to the manufacturer’s instruction. RT-PCR was performed using the same reagents and system as qPCR for MCMV DNA detection. TaqMan Gene Expression master mix was used for 18S (Mm03928990_g1), HPRT (Mm00446968_m1), IFIT1 (Mm00515153_m1), TNF (Mm00443258_m1), IL1β (Mm01336189_m1), IFNα (Mm00833976_s1), IFNβ1 (Mm00439552_m1), STAT2 (Mm00490880_m1), and SHH (Mm00436528_m1) (Life technologies, Foster City, CA). 18S or HPRT were used as internal control genes and fold changes for all experimental groups were expressed as 2^-ΔΔCt^ normalized to non-infected control group values.

### Laser microdissection (LMD)

PNd8 mice were sacrificed using CO_2_ inhalation and perfused with PBS as described above. Subsequently, brains were harvested and directly embedded in optimum cutting temperature (OCT) compound and stored at −80°C. 15μm brain sections were cut and adhered to PEN membrane-coated glass slides (Leica Microsystems Inc, Buffalo Grove, IL) and frozen at −80°C. Prior to LMD, slides were thawed at room temperature for 1min, fixed for 1min in 70% ethanol, washed in RNase-free water, and stained with 0.2% (w/v) cresyl violet for 30 seconds. Sections were subsequently washed in water and dehydrated by graded series of ethanol (70, 95, and 100%) for 1min each and air dried for 5 min. A Leica LMD6 microscope system (Leica Microsystems, Wetzler, Germany) was used to perform microdissection of the cerebellar EGL and dissected regions were collected in 0.5ml tube in GTC lysis buffer containing beta-mercaptoethanol. The collection tubes were centrifuged, vortexed repeatedly, and placed directly in dry ice or stored at −80°C. RNA was isolated with RNeasy Mini Kit (Qiagen, Germantown, MD) with on-column DNase digestion according to the manufacturer’s manual. cDNA was synthesized and RT-PCR was performed as described.

### *In vivo* deoxyuridine labeling

Developing postnatal mice were injected i.p. with 150mg/kg body weight of thymidine analogs of 5-bromo-2-deoxyuridine (BrdU) (Sigma Aldrich, cat# B9285, St. Louis, MO) or 5-Iodo-2-deoxyuridine (IdU) (Sigma Aldrich, cat# I7125, St. Louis, MO). BrdU was dissolved in sterile 0.9% saline and incubated at 55°C on dry heat block for 15 minutes to make 25mg/ml stock solution (Mandyam et al., 2007). Soluble BrdU was cooled to room temperature and aliquots were stored at −20°C. BrdU precipitates from thawed aliquots were redissolved by incubating in 55°C on dry heat block for 5-10 min and cooled to room temperature before injection in mice. IdU was dissolved in 0.2N NaOH/0.9% saline to make 70mg/ml stock solution, in which pH was adjusted to 9 by adding HCl, and aliquots were stored in −20°C. IdU was further diluted in 0.9% saline to make 10mg/ml working stock before injecting in mice.

#### Cumulative labeling

The lengths of the GCP cell cycle phases were determined using cumulative BrdU labeling (Florio et al., 2012; Nowakowski et al., 1989; Verslegers et al., 2013). Briefly, MCMV-infected mice and non-infected control mice received repeated injections of BrdU every 2 hours (hrs) for up to 24 hrs (0, 2, 4, 6, 8, 10, 12, 14, 16, 18, 20, 22, and 24 hrs). After the respective cumulative BrdU pulse labeling for each time point, pups were sacrificed at PNd8 (1, 1.5, 2, 4, 6, 8, 10, 12, 14, 16, 18, 20, 22, 24, and 26 hrs after the first BrdU injection) (Figure 4A). Following perfusion, brains were harvested, fixed with 4% paraformaldehyde (PFA), sectioned, and stained for BrdU, doublecortin (DCX), and phospho-histone H3 (Ser10) (pHH3) (see Table 1 for information on primary antibodies). DCX is selectively expressed in differentiated GCs in the iEGL and was used to identify and exclude the post-mitotic cells from the analyses (Takacs et al., 2008). The BrdU labeling index (BrdU LI) is defined as the proportion of BrdU^+^ cells per total number of cells in the oEGL, which increased over time, and alternatively expressed as the Growth Fraction(GF) (BrdU LI = Growth fraction (GF)) (Nowakowski et al., 1989). Mitotic cells in the oEGL were stained for pHH3 and mitotic labeling index (mitotic LI; pHH3 LI) was quantified to determine the duration of G2/M-phase (Contestabile et al., 2009; Hendzel et al., 1997; Lian et al., 2012; Takahashi et al., 1993). The calculations used for determination of the BrdU LI and mitotic LI (pHH3 LI) are shown below. Analysis of the cumulative BrdU experiment provided the total cell cycle length (T_C_) and each phase of the cell cycle (T_G_1, T_S_, and T_G2+M_) in GCPs in the oEGL of non-infected and MCMV-infected mice cerebella.

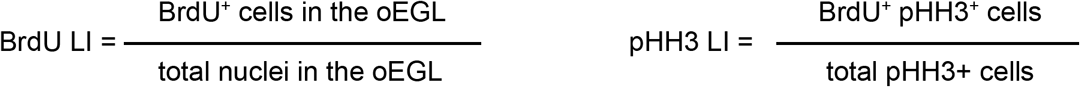

**Figure 1.**
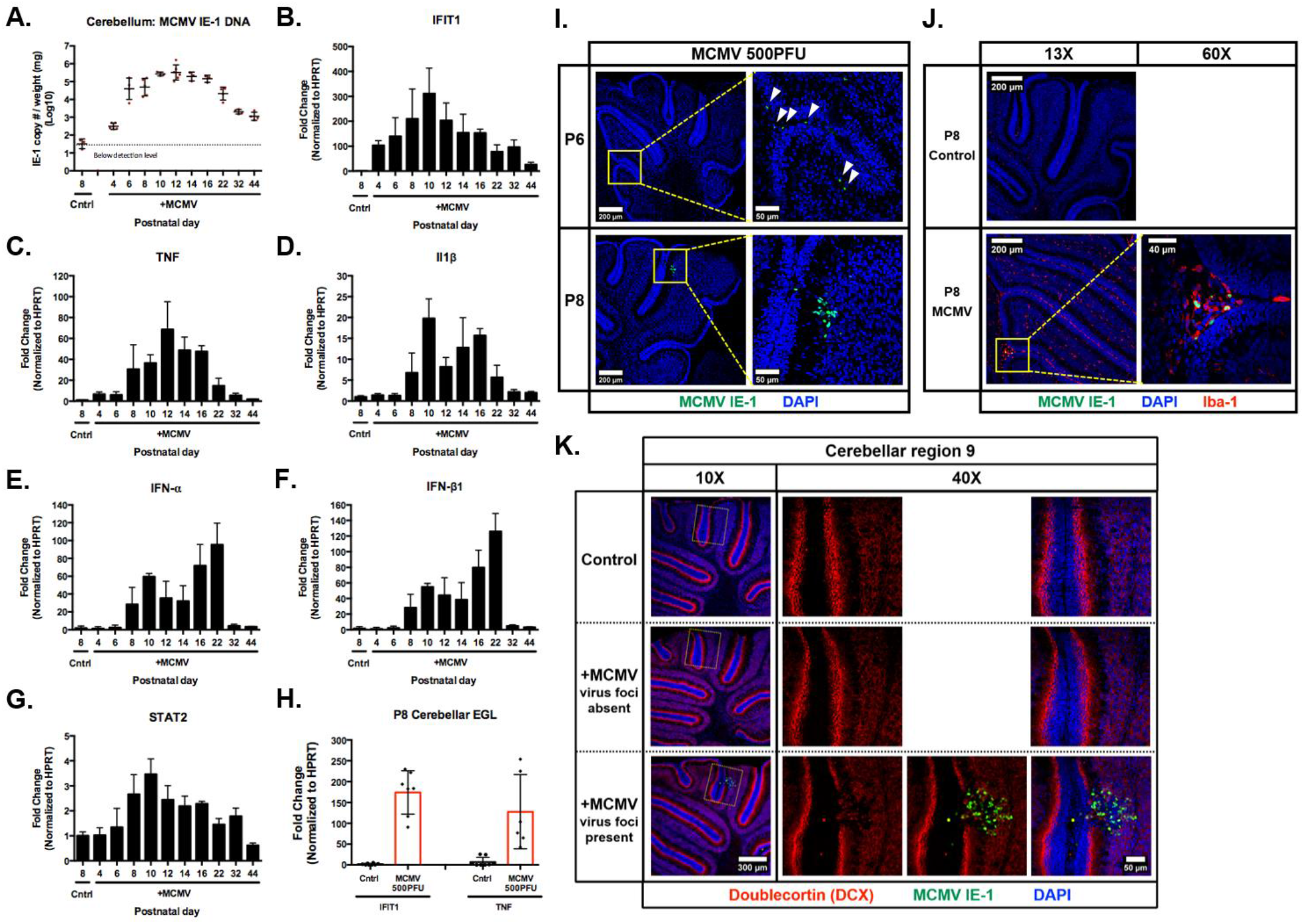
MCMV replicates in the cerebellum and induces a robust inflammatory response throughout postnatal period in newborn mice. Cerebella from MCMV-infected mice were homogenized and DNA/RNA extracted at various time points (PNd4-44) as described in the Materials and Methods. **(A)** Real-time PCR quantitation of MCMV DNA using total DNA extracted from the cerebellum. Each data point represents the genome copy number/mg of cerebellum. **(B-G)** Transcription of inflammatory mediators (IFIT1, TNF, IL1β, IFN-α, and IFN-β1) and transcription factor (STAT2) measured at different time in the postnatal period. HPRT was used as internal control to normalize the results and fold change was calculated by comparing RNA from cerebella from MCMV-infected animals to cerebella from non-infected, control animals. **(H)** Expression of IFIT1 and TNF were quantified by RT-PCR in RNA extracted from cerebellar EGL isolated by laser micro-dissection. **(I)** Distribution of MCMV infected cells in PNd6 (white triangles) and PNd8 cerebella detected with antibody reactive with MCMV IE-1 (pp89). Scale bar: 200 μm (left column) and 50 μm (right column). **(J)** Iba1+ mononuclear cells (red) in the cerebellum that also express MCMV IE-1 protein (green). Scale bar: 200 μm (13X images) and 40 μm (60X images). **(K)** PNd8 cerebellum double stained for MCMV IE-1 (green) and doublecortin (DCX) (red), which stains for immature/mature differentiated GCs. DAPI (blue) was used to stain the nucleus. Scale bar: 300 μm (10X images) and 50 μm (40X images). A total number of n=4-7 cerebella were used for each experiment.

**Figure 2.**
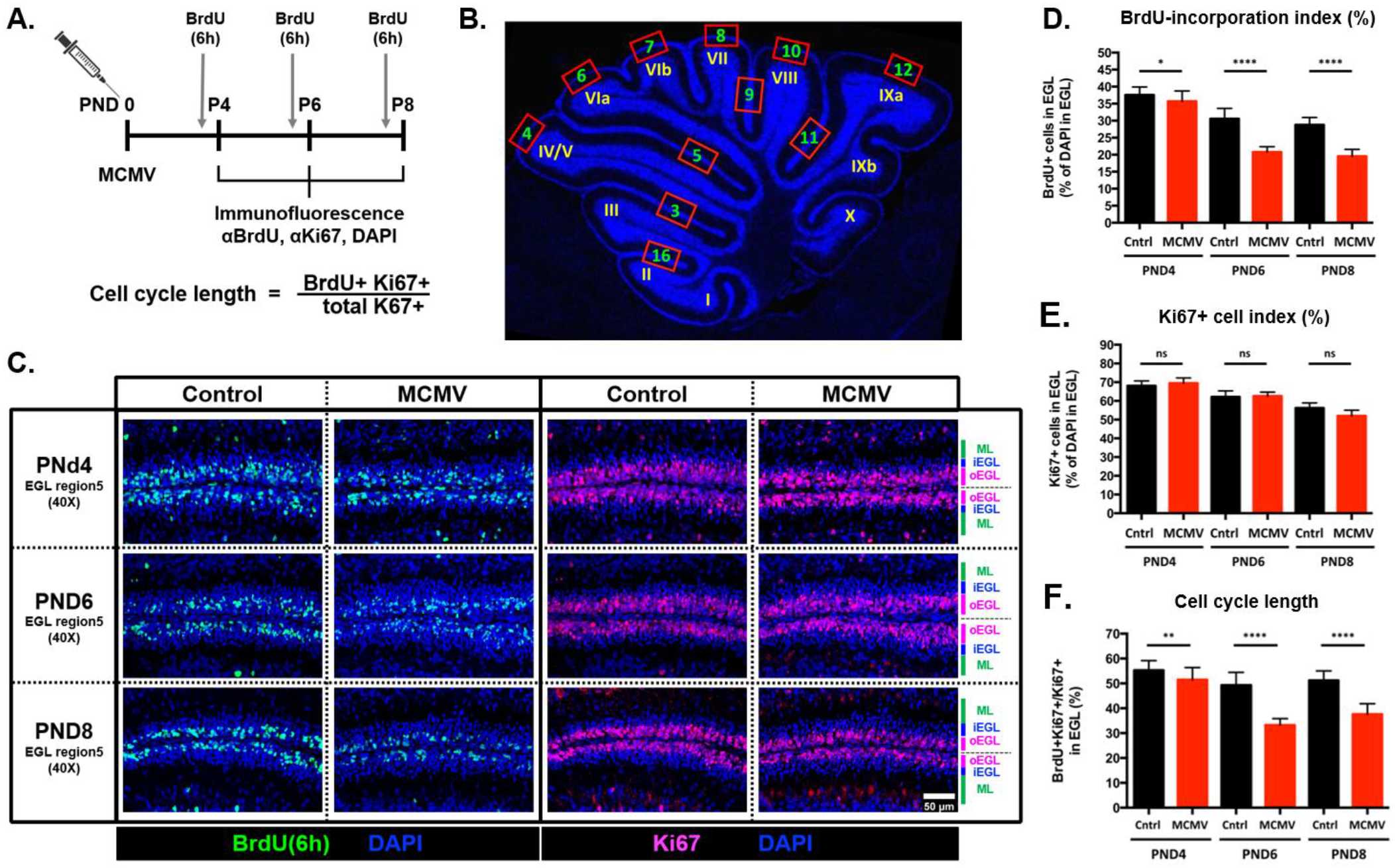
Cerebellar GCP proliferation is reduced and exhibit prolonged cell cycle length in MCMV-infected cerebella. **(A)** Schematic representation of the 6 hrs BrdU-incorporation experimental protocol and **(B)** designation of folia in cerebellum from PNd8 mouse. Analyses of the GCP quantifications were performed using cerebellar folia region 5. **(C-E)** Representative images and quantification of BrdU^+^(green) and Ki67^+^(magenta) GCPs at PNd4, 6, and 8 in the EGL of non-infected control and MCMV-infected mice cerebella. Scale bar: 50 μm. **(D)**The percentage of BrdU^+^ cells was decreased in the EGL of MCMV-infected mice at PNd6 and PNd8. **(E)** The percentage of Ki67^+^ cells from MCMV-infected mice from PNd6 and PNd8 were comparable to non-infected control mice. **(F)** Cell cycle length was estimated as percentage of Ki67 and BrdU double positive cells present in the population of Ki67^+^ cells (BrdU^+^Ki67^+^/total Ki67^+^ in the EGL (%)). A smaller percentage of double positive cells indicates a longer cell cycle. Data are shown as mean ± SD, n=4-6 mice/experimental group. P-values were calculated using two-tailed unpaired t-test (*p<0.05; **P<0.01; ****p<0.0001).

**Figure 3.**
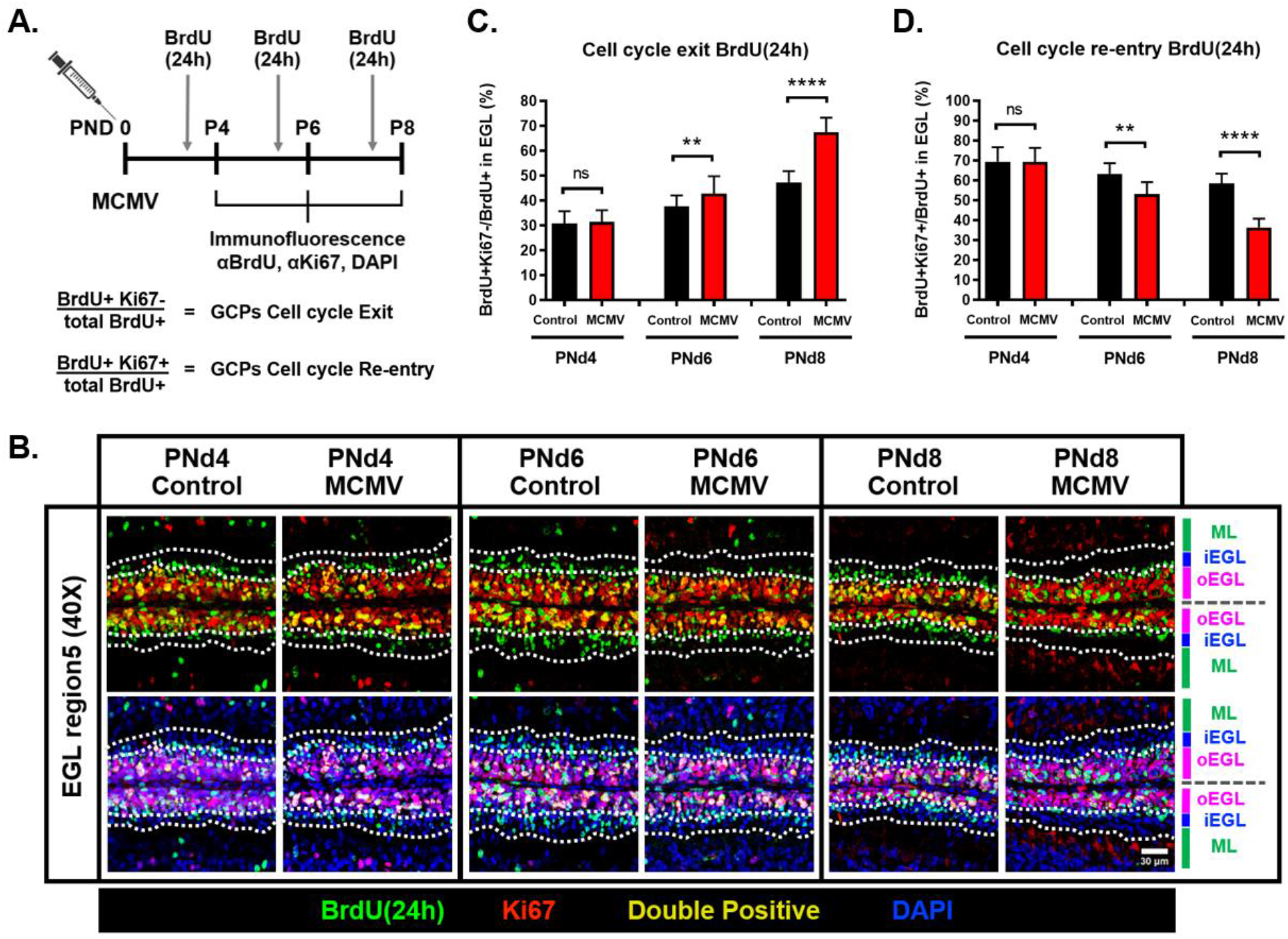
MCMV infection in newborn mice leads to increased cell cycle exit and delayed migration of GCPs from oEGL to iEGL. **(A)** Schematic representation of the experimental protocol for 24 hrs BrdU-incorporation. **(B)** Representative images demonstrating that the number of cells that exited cell cycle in the EGL (BrdU^+^Ki67^-^, green) was moderately increased at PNd6 and significantly increased at PNd8 in MCMV-infected mice. In addition, the number of cells that migrated from the oEGL (Ki67^+^ layer) to the iEGL (between two white dotted lines, Ki67^-^ layer) was reduced in cerebella from PNd6 and PNd8 MCMV-infected mice. Scale bar: 50 μm **(C-D)** Quantification of BrdU incorporation at PNd4, 6, and 8 as an estimate of cell cycle exit and re-entry. Brain sections were stained with anti-Ki67 and anti-BrdU antibodies 24 hrs after BrdU treatment. **(C)** Cell cycle exit was defined as ratio of the cells that were no longer in cell cycle as defined by the number of BrdU^+^Ki67^-^ cells to all cells labeled with BrdU (green) but not Ki67 (red) (GCP cell cycle exit = BrdU^+^Ki677^-^/total BrdU^+^ in the EGL (%)). **(D)** Similarly, cell cycle re-entry was defined as a ratio of cells that re-entered the following cell cycle as represented by ratio of the BrdU^+^Ki67^+^ cell population (yellow) to all cells labeled with BrdU(green) (GCP cell cycle re-entry = BrdU^+^Ki67^+^/total BrdU^+^ in the EGL (%)). Data are shown as mean ± SD, n=4-6 mice/experimental group in region 5 of the cerebellum. P-values were calculated using two-tailed t-test (*p<0.05; **P<0.01; ****P<0.0001).

**Figure 4.**
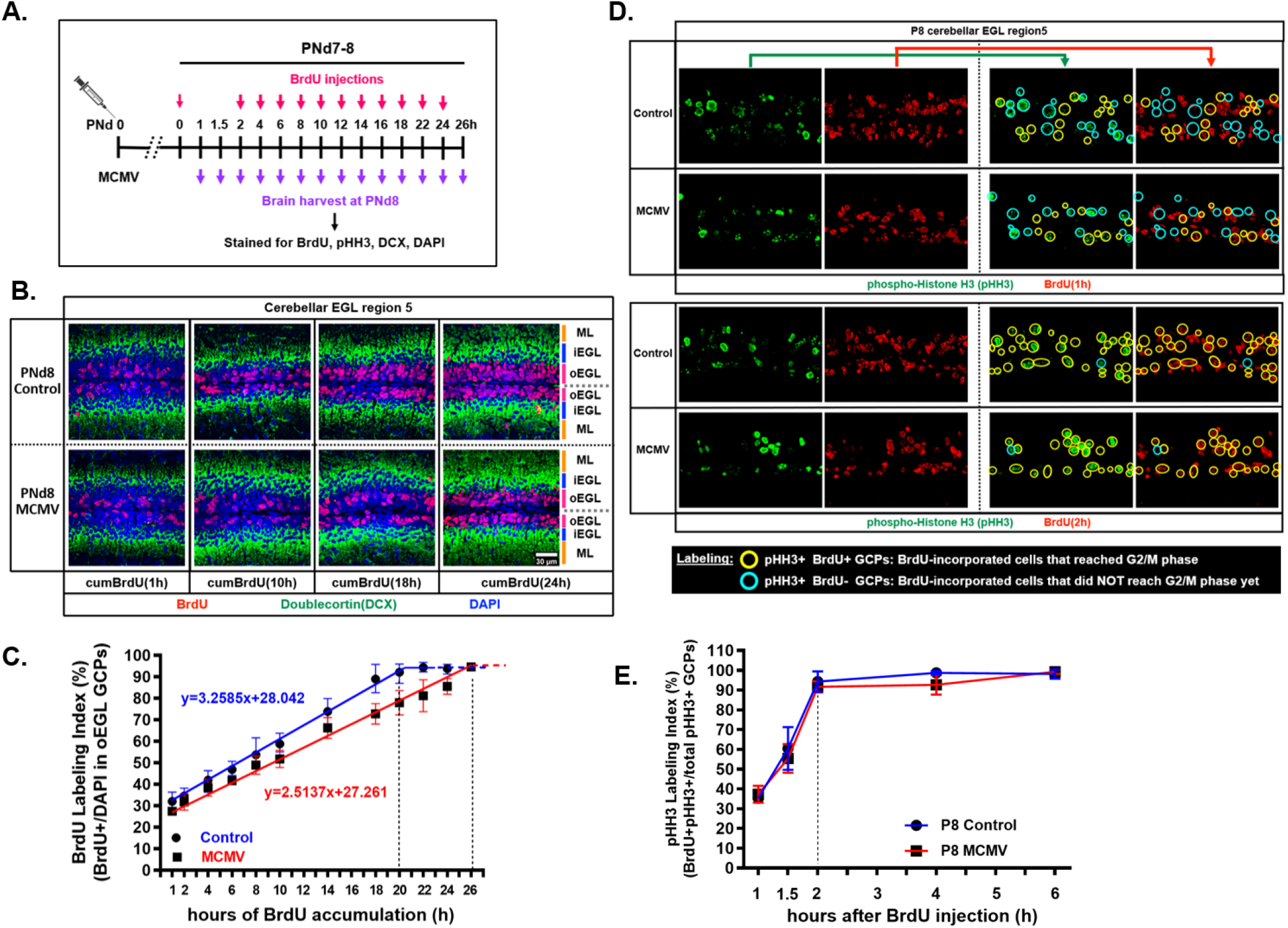
Prolonged GCP cell cycle length is due to the lengthening of G1- and S-phases but not G2/M-phase during MCMV infection. (**A**) Schedule of the *in vivo* cumulative BrdU labeling protocol to estimate the time required for each phase of the cell cycle of GCPs in the cerebellar EGL. **(B)** Representative confocal images of brain sections stained for BrdU (red), DCX (labeling the iEGL; green) and DAPI (nuclei staining; blue) from non-infected and MCMV-infected mice exposed to cumulative BrdU for the indicated time. In MCMV-infected cerebellum, fewer GCPs incorporated BrdU at all time points shown (1, 10, 18, and 24 hrs) compared to the non-infected, control mice. Scale bar: 30 μm. **(C)** BrdU^+^ GCPs in the oEGL of the cerebella were quantified (BrdU LI) at all time points and plotted against the duration of BrdU exposure to estimate cell cycle parameters. The BrdU LI acquired from the experiment was used to determine the duration of the total cell cycle (T_C_) and the time required to complete S-phase (T_S_). **(D)** The duration of G2/M-phase (T_G2+M_) was determined by single-dose BrdU labeling for 1, 1.5, and 2 hrs and stained for BrdU and pHH3. pHH3 immunolabeling defined cells in the G2/M-phase. pHH3^+^BrdU^-^ cells (light blue open circle) are pHH3^+^ cells that were not in S-phase (BrdU^-^) at the time of BrdU injection. BrdU^+^pHH3^+^ cells (yellow open circle) are cells that incorporated BrdU in the S-phase (BrdU^+^) and reached G2/M-phase. Scale bar: 50 μm. **(E)**pHH3 labeling index (pHH3 LI= BrdU^+^pHH3^+^/total pHH3^+^ GCPs in the EGL (%)) was determined. Data are shown as mean ± SD, n=4-6 mice/experimental group of the cerebellum. P-values were calculated using two-tailed t-test (*p<0.05; **P<0.01; ****p<0.0001).

**Table 1.** Primary antibodies used in this study.

| Antigen | Experiment | Company | Catalog # | Species | Isotype | Accession # |
| --- | --- | --- | --- | --- | --- | --- |
| IdU-FITC | IF | BD Bioscience | 347583 | Mouse | IgG1 | RRID:AB_400327 |
| BrdU [BU1/75(ICR1)] | IF | Abcam | ab6326 | Rat | IgG2a | RRID:AB_305426 |
| Ki-67 | IF | Abcam | ab66155 | Rabbit | polyclonal | RRID:AB_1140752 |
| Doublecortin (DCX) | IF | Abcam | 18723 | Rabbit | polyclonal | RRID:AB_732011 |
| Doublecortin (DCX) | IF | Santa cruz | sc-271390 | Mouse | IgG1 | RRID:AB_10610966 |
| TAG-1 (4D7) | IF | Iowa Hybridoma Bank |  | Mouse | IgM | RRID:AB_531775 |
| Calbindin-D28K | IF | Sigma | C9848 | Mouse | IgG1 | RRID:AB_476894 |
| MCMV IE-1 (pp89) | IF | Jonjic Lab generated | Croma 101 | Mouse | IgG1 |  |
| Iba1 | IF | Wako | 019-19741 | Rabbit | polyclonal | RRID:AB_839504 |
| phospho-histone H3 (ser10) | IF | Millipore/Upstate | 05-817 | Rabbit | polyclonal | RRID:AB_1587130 |
| Smoothened (Smo) (E-5) | WB | Santa cruz | sc-166685 | Mouse | IgG2a | RRID:AB_2239686 |
| Gli1 | WB | Abcam | ab49314 | Rabbit | polyclonal | RRID:AB_880198 |
| Gli2 (C-10) | WB | Santa cruz | sc-271786 | Mouse | IgG1 | RRID:AB_10708124 |
| phospho-Cyclin D1 | WB | Cell signaling | #2921S | Rabbit | polyclonal | RRID:AB_330139 |
| Cyclin D1 (92G2) | WB | Cell signaling | #2978S | Rabbit | polyclonal | RRID:AB_2259616 |
| CDK4 (DCS-35) | WB | Santa cruz | sc-23896 | Mouse | IgG1 | RRID:AB_627239 |
| CDk6 (B-10) | WB | Santa cruz | sc-7961 | Mouse | IgG1 | RRID:AB_627242 |
| Cyclin E1 (HE12) | WB | Cell signaling | #4129S | Mouse | IgG1 | RRID:AB_2071200 |
| CDK2 (D-12) | WB | Santa cruz | sc-6248 | Mouse | IgG1 | RRID:AB_627238 |
| E2F-1 (KH95) | WB | Santa cruz | sc-251 | Mouse | IgG2a | RRID:AB_627476 |
| phospho-Rb Ser780 | WB | Cell signaling | #9307 | Rabbit | polyclonal | RRID:AB_330015 |
| phospho-Rb Ser795 (B-4) | WB | Santa cruz | sc-514031 | Mouse | IgM |  |
| phospho-Rb Ser807/811 | WB | Cell signaling | #8516 | Rabbit | polyclonal | RRID:AB_11178658 |
| Rb (IF8) | WB | Cell signaling | #9313 | Rabbit | polyclonal | RRID:AB_1904119 |
| β-actin (C4) | WB | Millipore Sigma | MAB1501 | Mouse | IgG2b | RRID:AB_2223041 |

#### Dual Labeling

To confirm the T_C_ and the length of S-phase (T_S_) obtained from cumulative BrdU labeling experiment, we performed *in vivo* sequential IdU-BrdU dual-labeling of GCPs. To estimate T_C_, pups were given a single i.p. injection of IdU for 18, 22, 25, or 28 hrs followed by a single injection of BrdU 30 minutes prior to harvesting brains at PNd8 (Figure 5A). To determine the T_S_, PNd8 pups were given a single i.p. injection of IdU for 6 or 8 hrs followed by a single i.p. injection of BrdU 30 minutes prior to harvesting their brains at PNd8 (Figure 5D). Following perfusion and sectioning of fixed brain, brain sections were stained for IdU and BrdU and the formulas shown below were used to calculate T_C_ (Bouchard-Cannon et al., 2013; Brandt et al., 2012; Ryan et al., 2017) and T_S_ (Brandt et al., 2012; Harris et al., 2018; Iulianella et al., 2008; Martynoga et al., 2005; Quinn et al., 2007).

**Figure 5.**
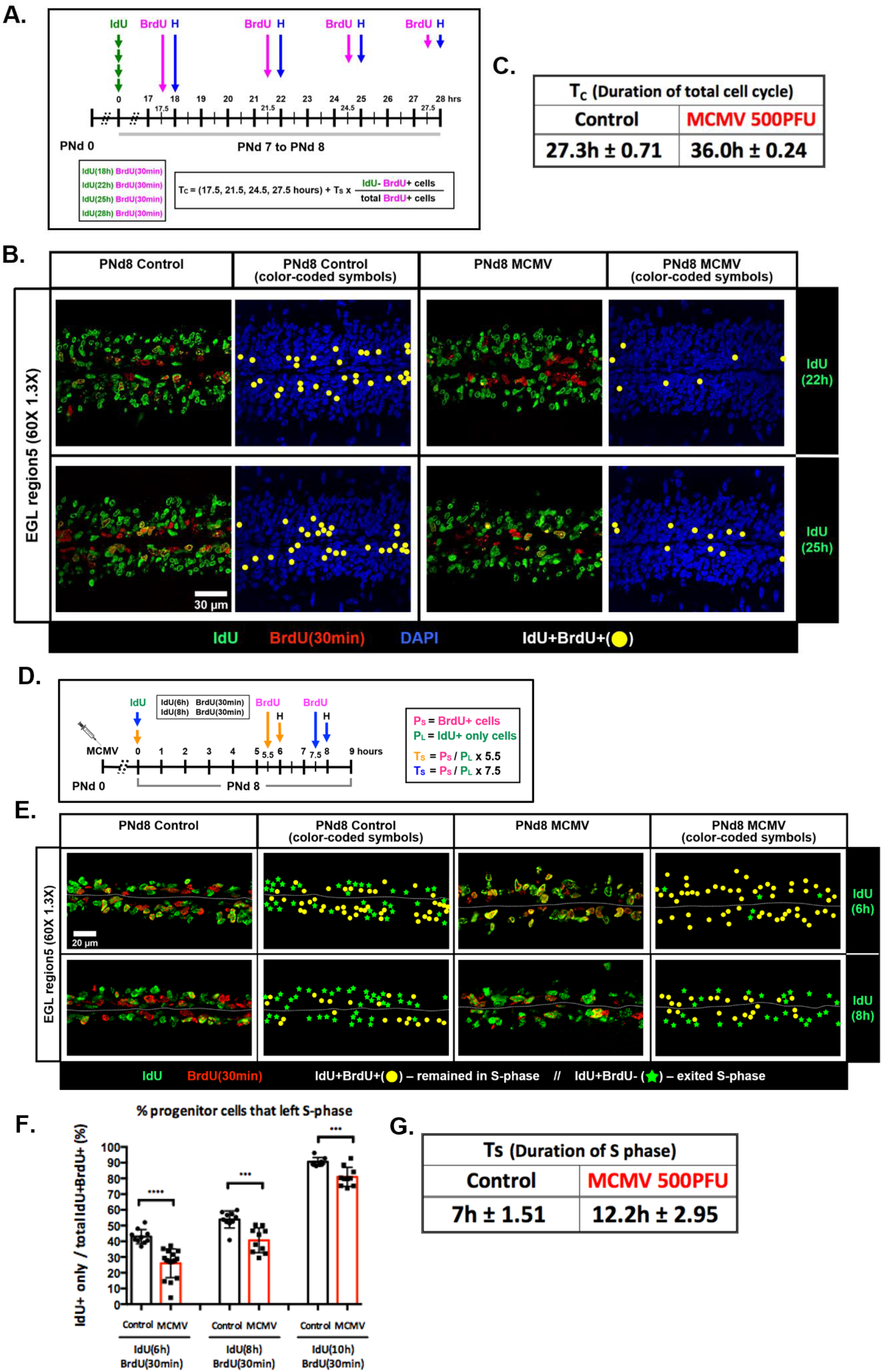
BrdU-IdU sequential labeling confirms the lengthening of the G1- and S-phases of the cell-cycle in MCMV-infected mice. **(A)** Schematic representation of the IdU-BrdU dual-labeling experimental protocol to measure the length of the total cell cycle (Tc). IdU was injected followed by BrdU injection at 17.5, 21.5, 24.5, and 27.5 hrs after the IdU injection. Brains were then harvested at PNd8. Cerebellar sections were stained for IdU (green), BrdU (red), and DAPI (blue) and IdU^+^ or BrdU^+^ GCPs quantified to analyze the T_C_ according to the equations indicated in the Material and Methods section. **(B)** Representative confocal images and images with color-coded symbols of cerebellar EGL indicate that reduced number of GCPs re-entered S-phase of the subsequent cell cycle (IdU^+^BrdU^+^ cells; yellow solid circle) in MCMV-infected mice compared to the non-infected, control mice during the time interval between IdU and BrdU injections (22 or 25 hrs). Scale bar: 30 μm. **(C)** T_C_ was approximately 8.7 hrs longer in GCPs in MCMV-infected mice cerebella compared to uninfected, control mice cerebella. **(D)**Schematic representation of the IdU-BrdU dual-labeling experimental protocol to measure the length of S-phase (T_S_). IdU was injected and BrdU was injected at 5.5 and 7.5 hrs after following the IdU injection and brains were harvested 30 mins after the BrdU injection on PNd8. **(E)** Representative images with color-coded symbols of cerebellar EGL indicate a reduced number of cells exited S-phase in the MCMV-infected cerebellum during the inter-injection intervals of 6 or 8 hrs. IdU^+^BrdU^+^ GCPs are population that remained in S-phase (yellow solid circle) and IdU^+^BrdU^-^ GCPs are population that left S-phase (green solid star). Scale bar: 20 μm. **(F)** Percentage of GCPs that left S-phase was quantified as IdU^+^/total IdU^+^BrdU^+^ GCPs in the EGL. **(G)** T_S_ was approximately 5.2 hrs longer in GCPs in MCMV-infected mice cerebella compared to uninfected, control mice cerebella. Data are shown as mean ± SD, n=4-6 mice/experimental group of the cerebellum. P-values were calculated using two-tailed t-test (*p<0.05; **P<0.01; ****p<0.0001).

T_C_ was determined by tracking GCPs that initially incorporated IdU and entered S-phase of the following cell cycle (IdU^+^BrdU^+^ cells) compared to GCPs that initially incorporated IdU but did not reenter the S-phase of the following cell cycle (IdU^+^BrdU^-^ cells) during the time interval between IdU and BrdU injections (Figure 5A). The T_C_ calculation was performed by quantifying cells that were in S-phase only during the second injection (IdU^-^BrdU^+^ cells) and total BrdU^+^ cells, which also included cells that were initially in S-phase that incorporated IdU and re-entered S-phase of the following cycle (IdU^+^BrdU^+^ cells) (Brandt et al., 2012) (Figure 5D). T_S_ was determined by quantifying GCPs that initially incorporated IdU and stayed in S-phase (IdU^+^BrdU^+^ cells) compared to GCPs that initially incorporated IdU but did not incorporate BrdU secondary to their exit from S-phase during the 6 or 8 hour interval between injections.

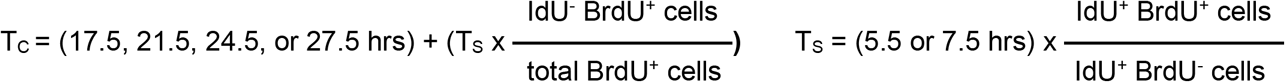

### Immunofluorescence and *in situ* hybridization

Animals were sacrificed by CO_2_ inhalation and perfused with PBS as described above. Brains were harvested and post-fixed in 4% PFA overnight at 4°C. Tissues were cryoprotected in 15% sucrose/PBS overnight followed by 30% sucrose/PBS for approximately 48 hrs at 4°C. Subsequently, cryoprotected brains were incubated in 30% sucrose/OCT (1:1) solution for 2-3 hrs at room temperature, embedded in optimal cutting temperature (OCT) medium (Electron Microscopy Sciences, Hatfield, PA), and snap frozen in 2-methylbutane/dry ice. Eight micron (8μm) brain sections were cut using a cryostat and sections were dried overnight at room temperature and either stained the day of processing or stored in −80°C.

For immunostaining, brain sections were rehydrated in PBS and incubated in blocking buffer (PBS, 0.3% TritonX-100, and 2% normal goat serum) for 3-4 hrs at room temperature followed by an overnight incubation with primary antibodies diluted in blocking buffer at 4°C. Primary antibodies (target, species, and source) used for immunofluorescence are listed in Table 1. Primary antibody incubation was followed by a 2-hour incubation in species/isotype-matched secondary antibodies conjugated with FITC, TRITC (SouthernBiotech, Birmingham, AL), or Alexa Fluor (ThermoFisher Scientific, Wiltham, MA) at room temperature. Hoechst dye (Hoechst 33342, trihydrochloride, trihydrate, Molecular probes, ThermoFisher Scientific, Wiltham, MA) was used to stain the nuclei. After immunostaining, brain sections were mounted with Vectashield (Vector Laboratories; Burlingame, CA) onto glass slides. For the detection of BrdU^+^ or IdU^+^cells, rehydrated brain sections were treated with 2N HCl for 20 minutes at 37°C and subsequently incubated in borate buffer (0.1M boric acid and 12.5mM sodium borate in deionized distilled water) for 10 minutes at room temperature to neutralize brain sections. Subsequent downstream procedures were identical to the immunostaining protocol described above.

Anti-TAG-1 immunostaining (Iowa Hybridoma bank, clone 4D7) utilized brain tissue from euthanized animals that were perfused with 1X PBS as described above followed by 4% PFA cardiac perfusion. This second perfusion was essential for TAG-1 immunostaining to visualize staining not only in the ridge (Figure 2B, regions 4,6,7,8,10,12) but also in the fissure (Figure 2B, regions 3,5,9,11,16) of the cerebellar folia. All other downstream procedures were identical to the immunofluorescence staining procedures described above. Fluorescence *In situ* hybridization (FISH) was performed as described previously (Morris et al., 2009). Probe constructs for *Gli1*, *Mycn*, and *Shh* were generated by Drs. A.L. Joyner (Memorial Sloan Cancer Center, NY), D.H. Rowitch (University of Cambridge, UK), and A.P. McMahon (University of California in San Francisco, CA), respectively. All three probes were generously provided to us by A.L. Joyner. Digoxigenin (DIG)-labeled RNA probes were transcribed *in vitro* using a DIG RNA labeling kit (Roche Applied Science, Basel, Switzerland) and brain sections were hybridized with gene-specific antisense RNA probes. Sense transcript was used as a negative control. *Shh* mRNA detection was coupled with protein staining for calbindin to visualize PCs. Detection of *Gli1* and *Mycn* mRNA were coupled with protein staining for DCX that stains the iEGL to localize *Gli1* and *Mycn* transcripts in the oEGL.

### Confocal imaging, cell counting and Statistical analysis

Cerebellar images were acquired using Olympus FV1000 confocal microscope and Olympus Fluoview FV10 ASW 4.2 software (Olympus, Tokyo, Japan). Identical magnification and laser power were used within the same experiment to acquire images of the cerebellar EGL. Subsequently, Fiji (ImageJ, NIH, Bethesda, MD) was used to process confocal images and to manually quantify GCPs by using the cell counter plugin (ImageJ > Plugins > Analyze > Cell counter). GCPs were quantified along the entire span of the EGL within the acquired images (164μm).

### GCP isolation

Primary GCPs were isolated from cerebella as described (Kenney and Rowitch, 2000; Lee et al., 2009). Briefly, four cerebella were pooled from PNd8 non-infected and MCMV-infected BALB/c mice and were dissected in calcium- and magnesium-free Hanks buffered saline solution (HBSS, Gibco/Life Technologies, Waltham, MA) supplemented with glucose. The meninges were stripped and cerebella were trypsinized (Worthington Biochemical Corp., Lakewood, NJ, cat# LS003736, 2.5% wt/vol) in 37°C for 15 minutes followed by trituration to make single cell suspension in HBSS/DNase/soybean trypsin inhibitor solution (Millipore-Sigma, Rockville, MD, cat# T2011, 1% wt/vol). Cells were filtered through a 70μm cell strainer and carefully placed on top of 4% Bovine Serum Albumin/HBSS and centrifuged at 70xg for 5min at 4°C to filter out large cells. To obtain a fraction enriched for GCPs, the cell suspension was loaded on a Percoll (GE Healthcare, Chicago, IL, cat#: 17-0891-01,) gradient of 35% and 65% and centrifuged at 1800×g for 10min at room temperature. GCPs were recovered from the 35%/65% interface and washed in HBSS/glucose solution. Recovered cells were pelleted and stored in −80°C until used for downstream experiments.

### Western blot

Samples from isolated GCPs were extracted by homogenization in RIPA buffer (50mM Tris-HCl pH7.5, 150mM NaCl, 1mM EDTA, 0.1% SDS, 1% NP-40, 0.5% Na-deoxycholate) containing Halt protease/phosphatase inhibitors (Thermo Fisher, Waltham, MA, cat# 78442). Homogenates were sonicated (10 sec sonicate/1min on ice/10 sec sonicate) and incubated on ice for 30 min-1 hour. Then samples were centrifuged at 13,200rpm for 10 min at 4°C. Total proteins were quantified using BCA assay (Pierce/ThermoFisher, Waltham, MA, cat# 23225) and equivalent (30μg) amounts of proteins per sample were loaded on a 9% SDS-polyacrylamide gel. The gel was transferred onto a 0.45μm nitrocellulose membrane (110V for 1hour 15min). Blotted membranes were blocked for at least 1 hour in 5% non-fat milk in TBS-t (10mM Tris-HCl, NaCl 150mM, pH 8.0)-Tween20 (0.1%) and incubated overnight with primary antibodies. Primary antibodies (target, species, and source) used for western blots are listed in Table 1. Membranes were washed and incubated at room temperature for 2-3 hrs with HRP-conjugated anti-rabbit, anti-mouse, or anti-rat secondary antibodies (SouthernBiotech, Birmingham, AL). The blots were developed using Western Lightning Plus-ECL (PerkinElmer, Waltham, MA) western blotting detection system. Densitometry was performed using Fiji (ImageJ, NIH, Bethesda, MD), and levels of proteins were normalized to β-actin. Immunoblotting utilizing some antibodies was carried out by horizontally cutting nitrocellulose membrane at a specific molecular weight to enable immunological detection of proteins that migrated markedly differently in an individual gel due to the difference in protein size. This approach was taken secondary to the limited amount of sample and the number of transferred gels required for the experiments. In addition, this enabled us to detect different targets from the same samples while limiting the background generated when antibodies from different species were applied to a single filter. Repeated stripping of nitrocellulose membrane was considered but pilot experiments provided unsatisfactory results secondary to residual background.

### Statistics

All statistical analyses were performed using Prism 6 (GraphPad, San Diego, CA). The Student’s t-test was used to compare statistical significance between two sample groups, non-infected and MCMV-infected mice. The Shapiro-Wilks test was used to analyze datasets for normality. Comparisons of multiple groups were subjected to ordinary one-way ANOVA with Tukey’s post-hoc multiple comparisons test for data with equal variances or otherwise by Dunn’s comparisons test to determine significance across treatment groups. Data are reported as mean ± standard deviation (SD). Values were considered to be statistically significant as indicated: (*) P < 0.05, (**) P < 0.01, (***) P < 0.001, (****) P < 0.0001, P-values above 0.05 (P > 0.05) was considered non-significant and is indicated as “ns”.

## Results

### MCMV replicates in the cerebellum and induces robust inflammatory response throughout postnatal development

Following i.p. inoculation of newborn mice with MCMV, MCMV DNA could be detected in the brain as early as PNd4, reaching a peak at PNd12, and then decreasing during subsequent time periods (Figure 1A). The expression of proinflammatory cytokines in MCMV-infected whole cerebella was quantified at these time points, including interferon-stimulating gene (IFIT1), type I interferons (IFN-α and IFN-β1), proinflammatory cytokines (TNF and IL-1β), and inflammation-associated transcription factor (STAT2). Expression of these markers of inflammation (e.g. IFIT1, TNF, IL1β, IFN-α, IFN-β1, and STAT2) followed similar kinetics of MCMV DNA throughout postnatal development (Figure. 1B-G). To further investigate potential sources of proinflammatory gene expression in the developing cerebellar cortex, we performed laser-capture microdissection (LMD) to isolate cerebellar EGL from PNd8 uninfected and MCMV-infected mice. Cerebellar EGL from infected mice exhibited robust increases in both IFIT1 and TNF expression, suggesting that cells within the EGL of infected mice contributed to the inflammatory response during viral infection even though this region of the cerebellum was not specifically targeted by the virus (Figure 1H). To further define the extent of MCMV infection in the cerebellum of newborn mice, a monoclonal antibody reactive against the major immediate early-1 (IE-1) protein of MCMV (pp89) was used to identify infected cells (Trgovcich et al., 2000). In the PNd6 cerebellum, MCMV infection was detected in single cells; however by PNd8, MCMV infection was identified as foci of infection in several different regions of the cerebellum as well as other regions of the brain including frontal cortex and hippocampus as previously described (Figure 1l) (Koontz et al., 2008). In addition, MCMV infection was occasionally detected in cells expressing Iba-1, a cellular marker for activated brain microglia or infiltrating monocytes (Figure 1J). Lastly, to determine the impact of foci of virus-infected cells on normal cerebellar morphology, PNd8 brain sections were stained for MCMV IE-1 and DCX, a microtubule-associated protein expressed in immature/mature differentiating and migrating GCs in the cerebellum (Gleeson et al., 1999). Areas of the cerebellum containing foci of virus infected cells displayed abberant morphology of DCX^+^ cells spanning the cerebellar cortex (Figure 1K). In contrast, minimal morphological alterations were present in regions of the cerebellum that did not contain foci of virus-infected cells. Together these results confirmed previous results that peripheral infection of newborn mice with MCMV led to productive virus infection in the cerebellum characterized by widely scattered foci of infection and the induction of a robust inflammatory response during postnatal cerebellar development (Koontz et al., 2008; Kosmac et al., 2013; Seleme et al., 2017).

### Cerebellar granule cell precursors (GCPs) exhibit prolonged cell cycle length and increased cell cycle exit in MCMV-infected cerebella

Despite the focal nature of MCMV infection in the brain, the deficits in the cerebellar cortical development were global and importantly, symmetric. We have previously shown that MCMV infection is associated with abnormal cerebellar cortical development including a decrease in cerebellar area and foliation, thinner IGL and ML, and a thicker EGL (Koontz et al., 2008). In earlier studies, we found a decrease in the BrdU incorporation of GCPs in the EGL, indicating that even though the EGL was thicker in infected mice, fewer cells were actively proliferating (Koontz et al., 2008). This previous finding was interpreted as evidence that in infected animals, the proliferation of GCPs in the EGL was delayed or perhaps incomplete and as a result, were also delayed in their subsequent differentiation from GCPs into GCs and migration from EGL to IGL. This mechanism was postulated to account for the thickened EGL in MCMV-infected mice. To investigate the possibility of altered GCP proliferation in MCMV-infected mice, we assessed cell cycle kinetics by measuring the cell cycle length, cell cycle exit, and cell cycle re-entry of the GCPs (Chenn and Walsh, 2002). First, GCP cell cycle progression was assayed in cerebella from MCMV-infected and non-infected mice to allow direct comparison of the relative length of the cell cycle. BrdU injections were given intraperitoneally to PNd4, 6, and 8 mice six hrs prior to harvesting brains, and brain sections were immunostained for BrdU (S-phase marker) and Ki67 (general proliferation marker), a protein that can be detected during all phases of the cell cycle (G1, S, G2, and M phases), except cells in G0-phase or cells exiting cell cycle (Figure 2A). The percentage of Ki67^+^ cells in cerebella from MCMV-infected mice were comparable to those in cerebella from non-infected, control mice at the three timepoints. (Figure 2C,E). Conversely, the percentage of BrdU^+^ cells was decreased in MCMV-infected cerebella as early as PNd4 with an even greater reduction in BrdU^+^ cells observed in cerebella from PNd6 and PNd8 mice (Figure 2C-D). The reduced proliferative capacity of GCPs in infected cerebella was also indicated by the decreased number of total mitotic cells as detected with an antibody reactive with phospho-histone H3 (pHH3) in the EGL of MCMV-infected mice (Supplemental Figure 1). The discrepancy in the results from two proliferation markers, BrdU and Ki67, suggested that there was a block or delay of cell cycle progression in GCPs in MCMV-infected mice. GCP cell cycle length was further estimated by calculating the labeling index as a percentage of BrdU^+^Ki67^+^ cells in total Ki67^+^ cells ((BrdU^+^Ki67^+^/total Ki67^+^) × 100 (%)). A smaller percentage would argue that the cell cycle is longer relative to GCPs from control mice. In MCMV-infected cerebella, cell cycle length of GCPs was slightly extended at PNd4 and was significantly prolonged at PNd6 and PNd8, as indicated by the decreased labeling index (Figure 2F). Together these data suggested that GCPs in the MCMV-infected mice were cycling more slowly than GCPs in non-infected, control mice and that lengthening of the cell cycle in GCPs in infected mice could contribute to an overall reduction in the number GCs.

We then determined if GCP cell cycle exit and re-entry was affected in MCMV-infected mice as compared to uninfected control mice. Control and MCMV-infected mice were pulse labeled with BrdU 24 hrs prior to harvesting brains at PNd4, 6, and 8, and fixed brain sections were stained and analyzed by immunofluorescence using antibodies against BrdU and Ki67 (Figure 3A). The percentage of cells that exited cell cycle was defined by the proportion of BrdU^+^Ki67^-^ cells in the total population of cells labeled with BrdU. Similarly, cell cycle re-entry was quantified by calculating the percentage of cells that re-entered cell cycle, defined as the proportion of BrdU^+^Ki67^+^ cell in the total population of cells labeled with BrdU during the 24-hour interval (Chenn and Walsh, 2002). Compared to uninfected controls, a significant increase in the percentage of GCPs that exited cell cycle as well as decreased number of cells re-entering cell cycle was observed in EGL from PNd6 and PNd8 but not in EGL from PNd4 MCMV-infected mice (Figure 3B-D). These observations were consistent with previous findings and provided additional evidence that a reduced number of GCPs progressed through the cell cycle, thus accounting decreased proliferation of GCPs and cerebellar hypoplasia that were observed in mice infected in the early postnatal period with MCMV.

After GCPs proliferate and exit cell cycle in the oEGL, GCPs differentiate into immature GCs, move into the deeper layer of the EGL, the iEGL, and then migrate radially along the Bergmann glial fibers to establish themselves in the IGL (Butts et al., 2014). In the absence of the initial migration from the oEGL to the iEGL, subsequent migration into the IGL is eliminated (Goldowitz and Hamre, 1998). We have previously reported that increased thickness and cellularity of the cerebellar EGL is associated with delayed radial migration of GCPs from EGL to IGL in MCMV-infected mice compared to the non-infected, control mice (Koontz et al., 2008). To further characterize the developmental abnormalities associated with altered proliferation and premature cell cycle exit of the GCPs in the EGL of MCMV-infected mice, we investigated the radial migration of the GCPs within the EGL of brain sections from mice that were pulse labeled with BrdU for 24 hrs. Immunofluorescence using anti-BrdU and anti-Ki67 antibodies revealed a significant reduction in the number of BrdU^+^ GCPs that migrated from oEGL to iEGL at PNd6 and PNd8, but not at PNd4 in MCMV-infected mice cerebella compared to cerebella from age-matched, non-infected control mice (Figure 3B). To confirm this finding, we performed a time course analysis with IdU, an alternative thymidine analog, by pulse-chase labeling for 18, 22, 25, 28, 32, 36, and 40 hrs in MCMV-infected and uninfected, control mice and harvested their brains at PNd8 to determine if migration of GCPs was prevented or delayed in MCMV-infected mice as detected by the presence of IdU^+^ GCPs in the iEGL. The iEGL was visualized by immunostaining with antibody against Transiently expressed Axonal Glycoprotein (TAG-1), a contact-related adhesion molecule that has been shown to have antagonistic effects in regulation of SHH-induced GCP proliferation (Bizzoca et al., 2003; Xenaki et al., 2011). We found maximum number of IdU^+^ cells in the iEGL at 32 hrs for GCs of non-infected, control mice and 40 hrs for GCs of MCMV-infected mice, suggesting that MCMV infection resulted in a significant delay in the initial migration of GCs from the oEGL to the iEGL (Supplemental Figure 2). Together these results argued that MCMV infection in newborn mice dysregulated GCP cell cycle progression, increased cell cycle exit, and delayed GC migration within the EGL of the cerebellar cortex.

### GCPs in MCMV-infected mice leads to increased cell cycle duration due to increase in G1- and S-phases during postnatal development

Previous studies have suggested that cell cycle kinetics are closely linked to cell cycle exit and neuronal differentiation (Schultze and Korr, 1981). Overexpression of cyclin D1/Cdk4 in NPCs in the murine developing cerebral cortex shortened the length of G1 and inhibited neurogenesis, whereas, lengthening of G1-phase by cyclin D1/Cdk4 inhibition promoted neurogenesis (Lange et al., 2009). These findings argue that increasing the duration of G1-phase of the cell cycle is sufficient to induce cell cycle exit of NPCs followed by their differentiation. Based on these findings, we utilized *in vivo* cumulative BrdU labeling to estimate total cell cycle length as well as length of different phases of the cell cycle of GCPs in non-infected and MCMV-infected mice. We administered BrdU to uninfected control and MCMV-infected mice every 2 hrs from 0 to 24 hrs and harvested brains at PNd8, as described previously (Nowakowski et al., 1989) (Figure 4A). Brain sections were stained with antibodies reactive with BrdU and a marker for differentiated GCs in the iEGL, DCX, in order to exclude post-mitotic cells from the analyses (Takacs et al., 2008). GCPs that were BrdU^+^ in the oEGL of the cerebella were quantified at each time point to generate a BrdU LI (ratio of BrdU^+^ GCPs to total cell number). The GCPs in the oEGL reached the maximum BrdU LI over time in both groups, in agreement with our findings that GCPs were cycling in infected and uninfected mice (Figure 4B-C). However, the time point at which the BrdU LI reached plateau was longer for GCPs in the MCMV-infected mice (26 hrs) compared to the non-infected, control mice (20 hrs) (Figure 4C). To obtain total cell cycle length (T_C_) and duration of S-phase (T_S_), the duration of BrdU exposure was plotted against BrdU LI and the best-fitted line was calculated. T_C_ and T_S_ were increased by 8.06 hrs and 2.02 hrs, respectively, in the GCPs of MCMV-infected mice (Table 2). In addition, to determine the duration of G2/M phase (T_G2+M_), PNd8 mice were given single-pulse BrdU injection and brains were harvested after 1, 1.5, and 2 hrs and the number of pHH3^+^ and BrdU^+^ GCPs was quantified. Mitotic LI was estimated by calculating the percentage of BrdU^+^pHH3^+^ cells of the total pHH3^+^ cells (BrdU^+^pHH3^+^/total pHH3^+^ cells) × 100 (%)).This allowed us to estimate the minimum time required for BrdU^+^ GCPs to enter G2/M (pHH3^+^). The maximum mitotic LI of GCPs was reached within 2 hrs in both non-infected, control and MCMV-infected cerebella (Fig 4D-E). The duration of G1-phase, derived by subtracting T_S+G2+M_ from T_C_, was increased by 6 hrs in GCPs from MCMV-infected mice as compared to the length of G1 in GCPs from uninfected control mice (Figure 4D, Table 2).

**Table 2.** Cell cycle parameters of GCPs from MCMV-infected and non-infected, control mice cerebella at PNd8 estimated by cumulative BrdU labeling.

| | $T_c-T_s$ (h) | $T_c$ (h) | $T_s$ (h) | $T_{G1}$ (h) | $T_{G2+M}$ (h) | Growth fraction |
| --- | --- | --- | --- | --- | --- | --- |
| **P8 Control** | **20** | **28.25** | **8.50** | **18** | **2** | **93.41% ± 1.65** |
| **P8 MCMV 500PFU** | **26** | **36.31** | **10.52** | **24** | **2** | **94.65% ± 1.70** |

The standard analysis of cumulative BrdU labeling experiment assumes that proliferating cells are asynchronous with cells uniformly distributed throughout the cell cycle and that the cell population increases in number with steady state dynamics as predicted from models of asymmetric cell division (Nowakowski and Hayes, 2008; Nowakowski et al., 1989). However, live cell imaging of *ex vivo* wholemount cerebella has provided evidence that at later stages of the developing cerebellum (PNd10), only approximately 4% of GCPs progress through asymmetric division while majority of GCPs go through terminal symmetric division (~73%) or non-terminal symmetric division (~22%) (Yang et al., 2015). Thus, to confirm the increased duration of T_C_ and T_S_ in GCPs of infected mice cerebella derived from the cumulative BrdU labeling studies, we utilized an assay based on sequential IdU-BrdU double-labeling (Figure 5). Results from cumulative BrdU labeling estimated that T_C_-T_S_ in control and MCMV-infected mice were 20 and 26 hrs, respectively (Table 2). Therefore, to determine T_C_ by sequential IdU-BrdU labeling, we chose an interinjection interval that encompassed the estimated T_C_-T_S_ determined by the cumulative BrdU labeling to ensure that we detected both populations of IdU^+^ cells that either entered the S-phase of the following cell cycle (IdU^+^BrdU^+^ cells) or did not reenter or exited the S-phase of the following cell cycle (IdU^+^BrdU^-^ cells) during the time interval between IdU and BrdU injections. Mice were pulse labeled with IdU for 18, 22, 25, or 28 hrs and injected with BrdU 30min prior to harvesting brains at PNd8 (Figure 5A). We observed reduced number of GCPs that entered the S-phase of the following cycle (IdU^+^BrdU^+^ cells) in the MCMV-infected mice (Figure 5B). The sequential IdU-BrdU labeling revealed that Tc was approximately 8.7 hrs longer in GCPs in MCMV-infected cerebella compared to uninfected, control cerebella, confirming the results from studies using cumulative BrdU labeling (Figure 5B-C). To measure Ts, PNd8 mice were pulse labeled with IdU for 6 or 8 hrs and injected with BrdU 30min prior to harvesting brains at PNd8 (Figure 5D). T_S_ was determined by counting the number of GCPs that remained in S-phase (IdU^+^BrdU^+^ cells) and compared to the number of cells that exited S-phase (IdU^+^BrdU^-^ cells) during the time interval between injections. GCPs from MCMV-infected mice displayed a significantly reduced number of cells that exited S-phase within 6 and 8 hrs and T_S_ was approximate 5.2 hrs longer relative to GCPs of non-infected control mice (Figure 5E-G). Together, these results indicate that in newborn mice, MCMV infection increased length of the cell cycle of GCPs from infected mice as a result of increased duration of G1- and S-phases but not G2/M-phase.

### GCP cell cycle signaling pathway is disrupted during MCMV infection

Sonic hedgehog (SHH) produced by Purkinje cells (PCs) is responsible for GCP proliferation in the cerebellar cortex during postnatal development (Dahmane and Ruiz i Altaba, 1999). To address whether reduced GCP proliferation was linked to altered SHH signaling in MCMV-infected cerebella, the expression of *Shh* mRNA within PCs was initially evaluated by *in situ* hybridization coupled with calbindin immunostaining to visualize PCs. *Shh* was readily detected in the PC bodies in non-infected, control cerebella whereas decreased levels of *Shh* mRNA expression were observed in MCMV-infected cerebella (Figure 6A). The decrease in *Shh* mRNA expression and SHH protein level were further validated by RT-PCR and western blotting of whole cerebella (Figure 6B-C).

**Figure 6.**
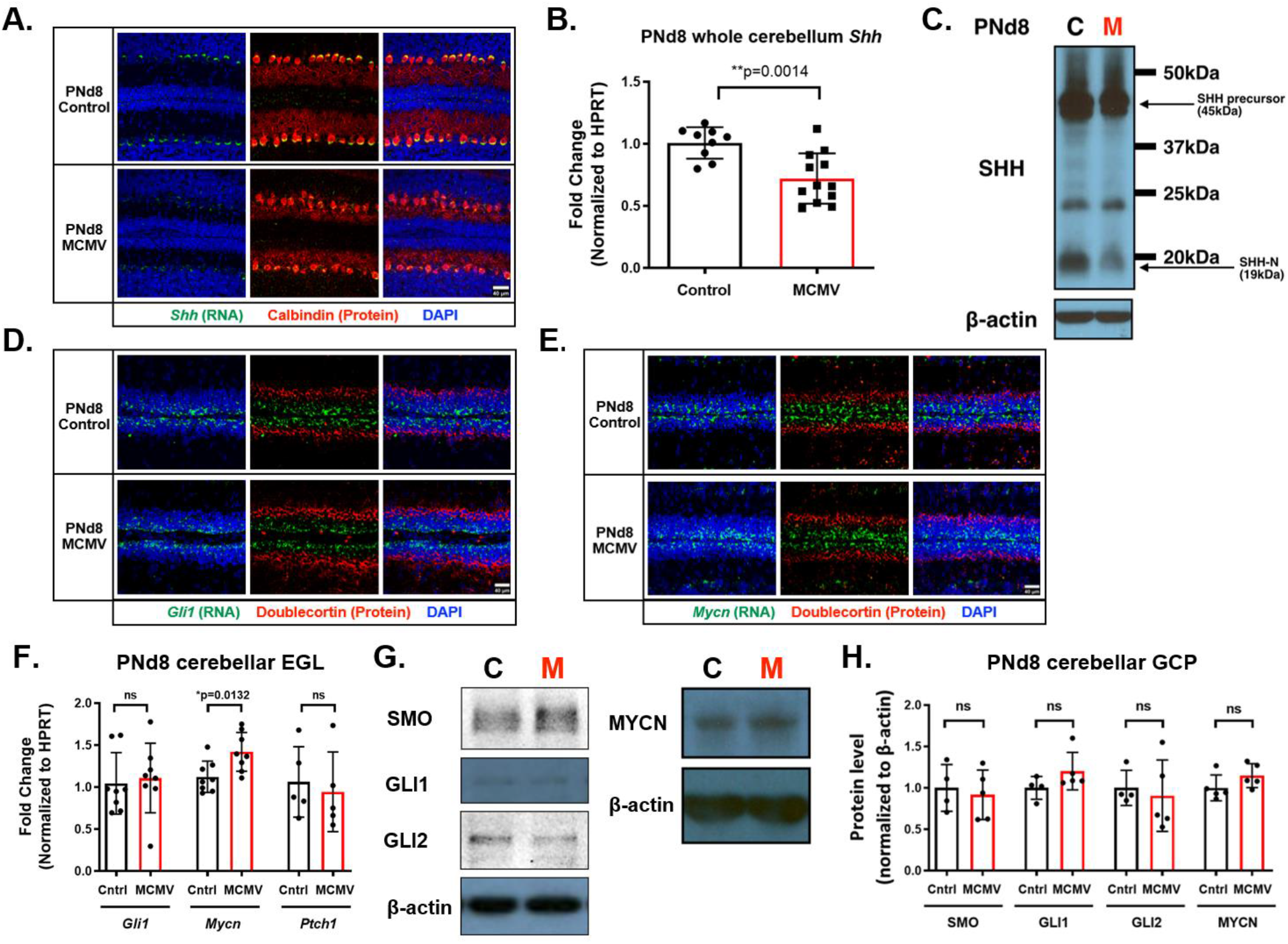
Sonic hedgehog (SHH) expression is decreased but SHH signaling pathways in GCPs of MCMV-infected cerebella are unaltered. **(A)** RNA *in situ* hybridization on PNd8 brain sections show reduced expression of *Shh* mRNA (green) in the Purkinje cells (red) detected with antibody against Calbindin in MCMV-infected mice. **(B)** RT-PCR and **(C)** western blotting of PNd8 whole cerebella confirmed the reduction of *Shh* mRNA expression and SHH protein level in the MCMV-infected mice. RNA *in situ* hybridization of **(D)** *Gli1* (green) and **(E)** *Mycn* (green), downstream transcription factors in SHH pathway were unaltered. Brain sections were immunostained for doublecortin (red) to label immature/mature differentiated GCs in the iEGL in the cerebellum. **(A,D,E)** Scale bar: 40 μm. **(F-H)** Genes and proteins downstream of SHH showed comparable levels between MCMV-infected and non-infected control cerebella. **(F)** *Gli1, Mycn,* and *Ptch1* transcript levels were measured from RNA isolated from laser micro-dissected cerebellar EGL. Note elevation of Mycn transcript in RNA from EGL. **(G-H)** Protein levels of SMO, GLI1, GLI2, and MYCN were measured and quantified from primary GCPs isolated from MCMV-infected and non-infected control cerebella. Data are shown as mean ± SD, n=4-6 mice/experimental group for immunofluorescence, n=8-12 mice/experimental group for RT-PCR, and 4-5 samples (4 cerebella pooled for each sample)/experimental group were used for western blot analysis. P-values were calculated using two-tailed t-test (*p<0.05; **P<0.01; ****p<0.0001).

SHH exerts its mitogenic effects on GCPs by binding to its receptor Patched1 (PTCH1) and alleviating PTCH1 repression of the G-protein coupled receptor Smoothened (SMO). SMO initiates a signaling cascade resulting in the increased expression of the transcription factors GLI and MYCN that upregulate target genes required for entry into G1-phase and cell proliferation (Kenney and Rowitch, 2000). To evaluate whether the reduction in SHH in PCs leads to concomitant changes in the SHH signaling pathway in GCPs, we determined the expression of key components downstream of *Shh*, *Gli1* and *Mycn*, by *in situ* hybridization Unexpectedly, similar levels of expression of *Gli1* and *Mycn* were detected in the EGL of cerebella from MCMV-infected and non-infected control mice (Figure 6D-E). These results were confirmed by RT-PCR of EGL samples obtained by laser microdissection of cerebella, which displayed similar or increased mRNA levels of G*li1*, *Ptch1*, and *Mycn* in the EGL of MCMV-infected animals compared to control animals (Figure 6F). In addition, comparable protein levels of SMO, GLI1, GLI2, and MYCN were detected in GCPs isolated from MCMV-infected and non-infected cerebella (Figure 6G-H). Collectively, these data demonstrated that while SHH production from PCs appeared to be decreased in MCMV-infected animals, the downstream effectors of the SHH signaling pathway in the GCPs appeared intact suggesting that altered SHH expression was not the primary factor affecting GCP proliferation, specifically cell cycle length, in infected animals.

We next examined the expression of proteins associated with the G1-phase and G1/S transition in GCPs isolated from PNd8 non-infected and MCMV-infected cerebella. The retinoblastoma (Rb) protein blocks S-phase entry by binding to E2F transcription factors that regulate progression through G1-phase and G1/S transition of the cell cycle. In mid G1-phase, cyclin D-Cdk4/6 complexes mono-phosphorylate Rb/p105 at residues including Ser780, Ser795, and Ser807/811 but Rb remains functional and continues to bind E2F transcription factors. During the late G1 restriction point, activation of cyclin E-Cdk2 complexes leads to Rb hyper-phosphorylation and release of E2F, promoting full E2F activity, and progression into S-phase (Boylan et al., 1999; Giacinti and Giordano, 2006; Narasimha et al., 2014). Thus, we examined the effect of MCMV infection on Rb, cyclin D1, and Cdk4/6 in the regulation of progression of G1-phase and G1/S transition (Delston and Harbour, 2006). While we observed unaltered levels of total Rb protein, we detected significantly reduced levels of Rb phosphorylated at Ser780 and Ser807/811 in GCPs from MCMV-infected cerebella relative to cerebella from uninfected control mice (Figure 7A-B). In contrast, the amount of Rb phosphorylated at Ser795 appeared similar in GCPs from either infected or non-infected mice (Figure 7A-B). The amount of E2F1 in MCMV-infected GCPs was comparable to that detected in GCPs from uninfected mice (Figure 7A-B). Levels of both cyclin D1 and p-cyclin D1 (Thr 286) were also unaltered; however, Cdk4/6 were differentially regulated as shown by a significant increase in Cdk4 and a significant decrease in Cdk6 protein levels in the MCMV-infected GCPs compared to the non-infected controls (Figure 7C-D). Lastly, mRNA expression and the amount of cyclin E as well as the amount of Cdk2 were decreased in GCPs of MCMV-infected mice as compared to cerebella from control mice (Figure 7C-E). Reduced levels of cyclin E and Cdk2 in infected animals could potentially limit hyper-phosphorylation of Rb, thus slowing the release of E2F and delaying the G1/S transition. Together, these data are consistent with the lengthened G1- and S-phases observed in GCPs from MCMV-infected mice when compared to control mice.

**Figure 7.**
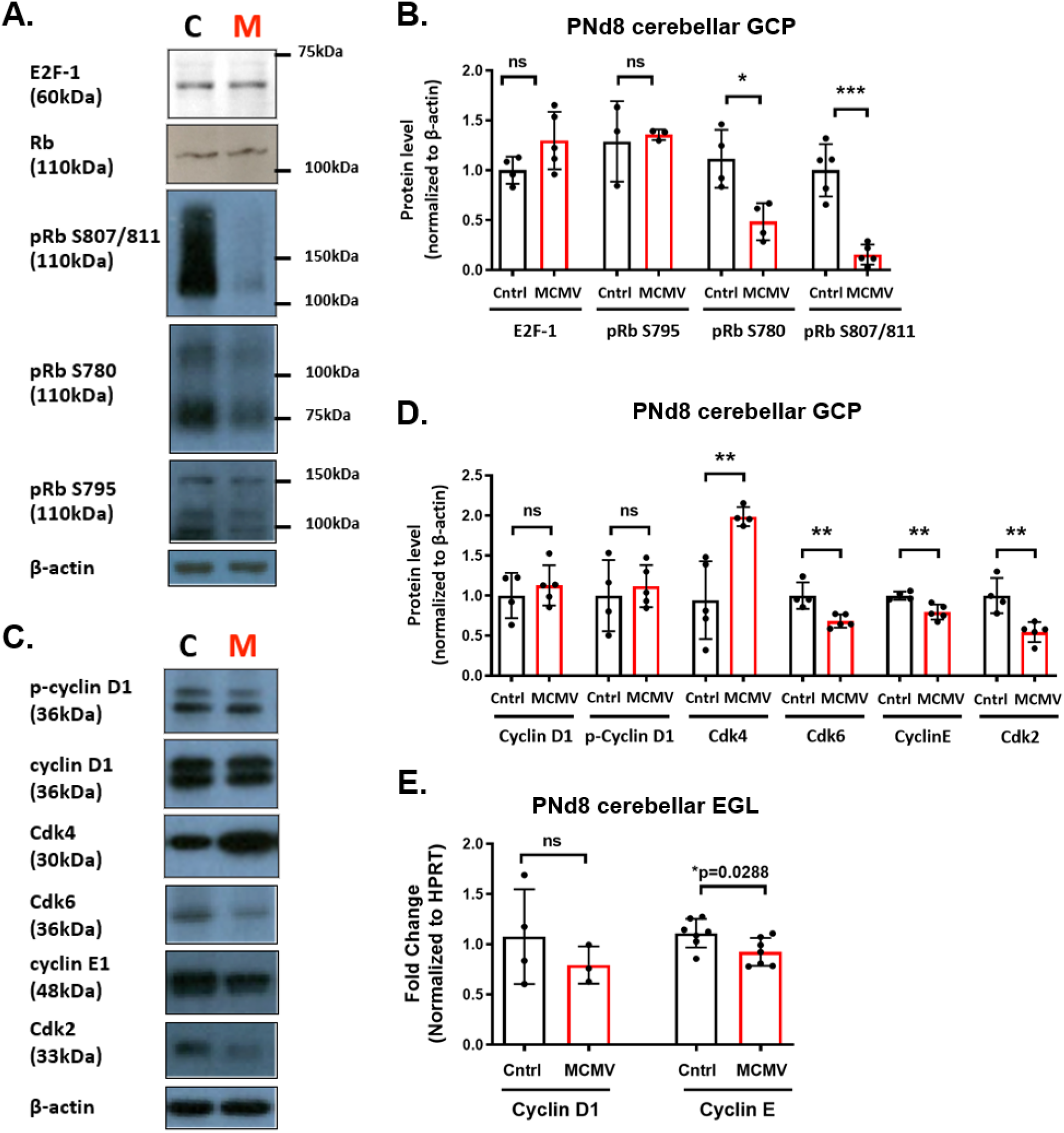
MCMV infection in newborn mice alters phosphorylation status of Rb and reduces protein levels of cyclin E and Cdk2. **(A-B)** The expression of cell cycle proteins was quantified by immunoblotting of GCPs isolated from PNd8 non-infected control and MCMV-infected cerebella. Protein expression of Rb and E2F-1 were quantified using *β-actin as an internal control.* Levels of E2F-1, Rb, and pRb S795 levels were unaltered; however pRb S780 and pRb S807/811 were significantly reduced in GCPs from MCMV-infected mice compared to non-infected, control mice. ***(C-D)** GCPs from MCMV-infected cerebella showed that Cdk4 and Cdk6 were differentially regulated while cyclin D1 or p-cyclin D1 were unaltered. Cyclin E and Cdk2 were reduced in GCPs from MCMV-infected mice cerebella. **(E)** Transcript levels of cyclin D1 and cyclin E from laser micro-dissected cerebellar EGL correlated with the protein levels measured from isolated GCPs*. Data are shown as mean ± SD. n=4-8 mice/experimental group for RT-PCR, and 4-5 samples (4 cerebella pooled for each sample)/experimental group were used for western blot analysis. P-values were calculated using two-tailed t-test (*p<0.05; **P<0.01; ****p<0.0001).

### Treatment with TNF-NAb normalizes the prolonged G1- and S-phases in GCPs of MCMV-infected mice

Previously we have shown that treatment of MCMV-infected newborn mice with either the anti-inflammatory corticosteroid (e.g. methylprednisolone) or TNF-Nab normalized many of the morphometric abnormalities in the cerebella of infected mice, suggesting that host inflammatory response to MCMV infection is, in part, responsible for the abnormal cerebellar development in mice infected with MCMV in the perinatal period (Kosmac et al., 2013; Seleme et al., 2017). Using a similar approach, we determined if TNF-NAb could normalize the cell cycle abnormalities that we had defined in infected mice. Infected and uninfected newborn mice were treated with vehicle, isotype control antibody, or TNF-NAb on PNd3-7 once daily and the impact on the cell cycle in GCPs was determined on PNd8 (Figure 8A). To estimate the impact of TNF-NAb treatment on the duration of the S-phase in infected animals, mice from all experimental groups were injected with IdU for 6 hrs followed by injection with BrdU 30min prior to harvesting brains (Figure 8A). Consistent with previous studies, altered GCP proliferation, characterized by decreased BrdU incorporation, was observed in the cerebella of vehicle-treated or isotype control antibody-treated mice following MCMV infection (Figure 2D, Figure 8B). Treatment of infected mice with TNF-NAb normalized the BrdU incorporation to the level comparable to uninfected, control mice (Figure 8B). The percentage of Ki67^+^ GCPs remained unchanged in all three treatment groups of MCMV-infected mice compared to uninfected, control mice (Figure 2E, Figure 8C) (Koontz et al., 2008; Kosmac et al., 2013; Seleme et al., 2017). In addition, the extended length of S-phase in GCPs of MCMV-infected mice that received vehicle or isotype control antibody was observed compared to non-infected, control mice, and this extended length of S-phase was normalized upon TNF-NAb treatment of MCMV-infected mice (Figure 8F).

**Figure 8.**
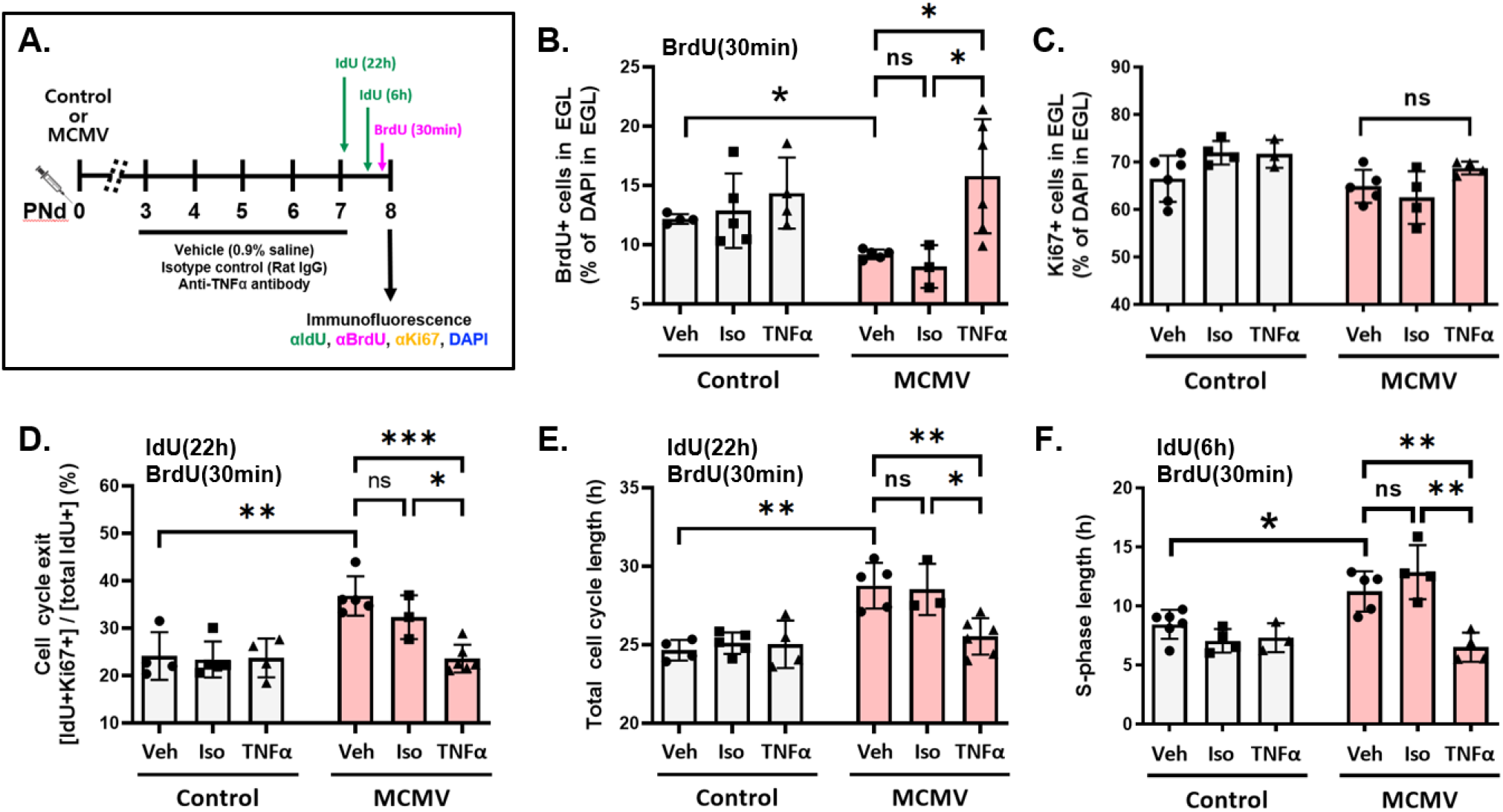
Treatment with TNF-NAb normalizes cell cycle abnormalities following MCMV infection. **(A)** Schematic representation of the IdU-BrdU dual-labeling experimental protocol in MCMV-infected mice and non-infected, control mice treated with vehicle (Veh), isotype control antibody (Iso), or TNF*α neutralizing antibody* (TNF-Nab) as described in Materials and Methods section. Harvested and fixed brain sections were stained for IdU, BrdU, Ki67, and DAPI to measure parameters of the cell-cycle as described in previous figures. **(B-C)** Brain sections from animals treated with BrdU for 30 min were analyzed to measure cell proliferation by quantifying **(B)** BrdU^+^ and **(C)** Ki67^+^ GCPs in the EGL. **(D-E)** Mice were pulse labeled with IdU for 22 hrs and injected with BrdU 30min prior to harvesting brains at PNd8 to estimate **(D)** cell cycle exit and **(E)** total cell cycle length. **(F)** Mice were labeled with IdU for 6 hrs followed by BrdU injection 30min prior to harvesting brains at PNd8 to estimate the length of S-phase of GCPs in the EGL. Data are shown as mean ± SD, n=4-6 mice/experimental group for immunofluorescence. P-values were calculated by using one-way ANOVA with Tukey’s multiple comparison’s test (*p<0.05; **P<0.01; ****p<0.0001).

To determine the proportion of cells exiting cell cycle and total cell cycle length, mice were treated with IdU for 22 hrs followed by BrdU injection 30 minutes prior to harvesting brains at PNd8 (Figure 8A). Treatment with TNF-NAb in MCMV-infected mice normalized the proportion of cells exiting cell cycle as well as the total cell cycle length of GCPs, while isotype control antibody treatment groups had no significant effect on the cell cycle regulation (Figure 8D-E). Overall, these data indicate that TNFα-regulated inflammatory responses induced by MCMV contributed to altered GCP proliferation in the cerebellar cortex of MCMV-infected mice.

## Discussion

Newborn infants infected *in utero* with HCMV can exhibit a variety of structural brain abnormalities, including cerebellar hypoplasia (Cheeran et al., 2009; de Vries et al., 2004; Manara et al., 2011; Tsutsui et al., 1997). To define mechanisms of CNS damage associated with intrauterine HCMV infection, we developed a murine model utilizing MCMV infection of newborn mice that recapitulates many of the findings observed in infants with congenital HCMV infection. Histopathological studies in this model have shown global and symmetric dysmorphogenesis of the cerebella of MCMV-infected mice with foci of virus-infected cells widely dispersed in different regions of the brain, including the cerebellum. Cerebellar hypoplasia, including reduced cerebellar size, area, and foliation, is a prominent feature of MCMV-infected animals and as noted above, is also observed in congenitally infected infants (Cheeran et al., 2009; Koontz et al., 2008). These abnormalities in infected mice are also accompanied by changes in the cortical structures of the cerebellum such as increased thickness of the EGL and decreased thickness of ML and IGL (Koontz et al., 2008). Animal models mimicking various examples of altered brain development (genetic or infectious agents) have identified several mechanisms underlying microcephaly including an increased number of apoptotic NPCs, reduced proliferation, or premature differentiation of NPCs (Barbelanne and Tsang, 2014; Cremisi et al., 2003; Miyata et al., 2010; Oh et al., 2017; Roper et al., 2006). In our previous studies of MCMV infection in newborn mice, we did not detect increased number of apoptotic cells in the infected cerebella compared to the control cerebella indicating that increased cell death was not responsible for reduced cerebellar size; however, we did observe reduced proliferation and delayed migration of GCPs in the cerebellar cortex (Koontz et al., 2008; Kosmac et al., 2013; Seleme et al., 2017). In the current study, we extended these observations to further define mechanism that contributed to altered cerebellar morphogenesis.

We measured the cell cycle length of GCPs in PNd8 MCMV-infected and non-infected control mice and determined that the cell cycle length of GCPs in the infected mice is longer, but importantly, the cell cycle is not arrested. Relative to the control samples, we observed a decreased number of BrdU^+^ GCPs while the number of Ki67^+^ GCPs remained unaltered in the cerebellar EGL of infected mice, indicating that while cells entered the cell cycle, there was a delay in progression of the cell cycle to S-phase (Figure 2F). Extension of these studies to include measurements of the length of G1-, S-, and G2/M-phases of the cell cycle revealed that the reduced proliferation of GCPs in infected mice could be attributed to the lengthening of both G1- and S-phases. Several explanations could account for this finding including direct viral damage to the GCPs or virus-induced host inflammatory response that alter GCP proliferation. Although appealing, a mechanism involving direct viral damage is a less plausible explanation because virus-infected cells appeared only in foci that were widely scattered throughout the parenchyma of the cerebellum, a finding inconsistent with the symmetric and global dysmorphogenesis of the cerebellum observed in this model. Moreover, we have rarely observed foci of infected cells in the EGL. For this reason, we believe that the prolonged cell cycle length of GCPs is more likely due to virus-induced inflammatory responses in the CNS. The interplay between virus-induced inflammatory responses and brain development was demonstrated in our previous studies in which either corticosteroid or TNF-NAb treatment decreased inflammation and limited abnormal CNS development including GCP proliferation deficits (Kosmac et al., 2013; Seleme et al., 2017). We have confirmed these earlier results in the current study and further refined our observations on the impact of blocking TNFα in infected mice by demonstrating that inhibiting TNF signaling normalized the cell cycle abnormalities in GCPs of infected mice (Figure 8). One potential mechanism to explain prolonged S-phase in GCPs of MCMV-infected mice is that inflammation can lead to DNA damage response (DDR) pathways. Prior studies have shown that the induction of DNA damage slows the rate of S-phase progression (Willis and Rhind, 2009). Thus in our mouse model of MCMV infection it is possible that the cell cycle of GCPs is slowing in S-phase in order to repair DNA damage incurred within the inflammatory environment. In particular, TNFα has been implicated to indirectly cause DDR through the production of ROS by altering the mitochondrial function (Schulze-Osthoff et al., 1992; Yan et al., 2006). Studies using TNFα stimulated L929 fibroblasts have provided evidence that TNFα induces mitochondrial ROS that contribute to cytotoxicity (Goossens et al., 1995; Shoji et al., 1995). Our analyses of cell proliferation were performed at PNd8, a time point at which there is an approximately 30-fold increase in the TNFα transcript level, which continues to increase through PNd12 in the cerebellum of MCMV-infected mice (Figure 1C). Therefore, one possible mechanism is that TNFα may induce mitochondrial ROS in the GCPs of MCMV-infected mice leading to genomic instability. However, we observed only rare cleaved caspase-3 positive GCPs in the cerebella of MCMV-infected mice, thus it seems unlikely that GCPs had significant DNA damage present in GCPs of infected mice. More recently, we have reported that interferon gamma (IFNγ) also contributed to the phenotype(s) of cerebellar abnormalities that we previously described in MCMV-infected newborn mice, including increased thickness of the EGL as compared to control animals (Kvestak et al., 2021). Furthermore, infiltrating NK/ILC1 produced IFNγ in the cerebella of MCMV-infected mice and were shown to contribute to the phenotype of increased thickness of the EGL of the cerebellum that we described in this model (Kvestak et al., 2021). In contrast to findings of the current study, the findings from the study of Kvestak, *et.al.*, (2021) were interpreted as evidence that IFNγ-driven expression of SHH lead to the increased thickness of the cerebellar EGL (Kvestak et al., 2021). Although results in the current study failed to demonstrate an increase in the expression of SHH in the cerebella of MCMV infected mice, there appeared to be some evidence of increased transcription of at least on downstream target of *Shh*, *Mycn*. However it is important to note that in the current study we quantified transcription of downstream targets of *Shh* in cells from the EGL isolated by laser microdissection and not from whole cerebellum. Thus, the importance of dysregulation in the SHH signaling pathway and observed thickened EGL phenotype in infected mice in this model remains to be further defined. However, together the findings in the current report and those in this previous study strongly argue that a combination of proinflammatory signaling molecules contribute to the phenotype of altered cerebellar development in this model of virus-induced inflammation during neurodevelopment. Lastly, we are cognizant that although our observations have provided additional understanding of the impact of virus-induced inflammation on neurodevelopment, additional studies will be required for a more precise definition of the mechanism(s) leading to altered neurodevelopment in this model.

GCP proliferation in the cerebellar cortex is widely known to be primarily regulated by SHH signaling (Dahmane and Ruiz i Altaba, 1999; Wallace, 1999; Wechsler-Reya and Scott, 1999). The soluble form of SHH is produced by the PCs of the cerebellum after binding to its receptor PTCH-1 on GCPs, triggers a signaling pathway that upregulates the expression of the GLI family of transcription factors and MYCN (Kenney et al., 2003). Surprisingly, while we detected decreased levels of SHH transcripts and protein consistent with alterations in the proliferative program of the GCPs of infected animals, the downstream SHH signaling pathway within the GCPs appeared intact (Figure 6). It is possible that the reduction of SHH in the cerebella of infected mice slowed the cell cycle progression while maintaining downstream SHH signaling and supporting GCP proliferation. Alternatively, it is also possible that the magnitude of the observed decrease in SHH expression was insufficient to alter downstream signal amplification. Lastly, insulin-like growth factor (IGF) has also been implicated as a potent mitogen for GCPs during postnatal development and synergizes with the downstream signaling pathway of SHH on GCPs (Fernandez et al., 2010). Although this remains a potential explanation for our finding, our previous transcriptomic studies indicated there was not a significant change in IGF expression in cerebella from infected mice as compared to uninfected control mice (Koontz et al., 2008).

Interestingly, despite the intact SHH signaling pathway in the GCPs of the infected mice, cell cycle regulatory proteins were altered, including the phosphorylation status of Rb and the expression of cyclins and Cdks responsible for G1-phase and G1/S transition. Cell cycle progression during mid-G1 phase first requires mono-phosphorylation of Rb/p105 at residues including Ser780, Ser795, or Ser807/811, by cyclin D-Cdk4/6 complexes (Boylan et al., 1999; Narasimha et al., 2014). Following the initial mono-phosphorylation of Rb, cyclin E-Cdk2 kinase activity leads to hyper-phosphorylation of Rb resulting in release of E2F transcription factors that regulate the expression of genes required for S-phase entry and DNA replication (Harbour et al., 1999; Koff et al., 1992). Since the kinase activity of the Cdks depends on binding to cyclins, the phosphorylation state of Rb can be viewed as an indirect measure of cyclin-associated kinase activity. In our studies, Rb phosphorylation at Ser795 was similar in MCMV-infected and control mice; however, phosphorylation at Ser780 and Ser807/811 was significantly reduced in GCPs of infected mice, raising the possibility that additional phosphorylation events and ultimately, the release of E2F transcription factors likely occured with reduced efficiency. This hypothesis is also consistent with our observation of decreased cyclin E and Cdk2 protein expression in the GCPs of infected mice (Figure 7C-D). We propose that decreases in cyclin E and Cdk2 protein levels lead to decreased kinase activity of the cyclin E-Cdk2 complexes, which subsequently reduced Rb hyper-phosphorylation and the expression of E2F-regulated genes essential for the G1/S transition. This interpretation of our findings would be consistent with lengthening of the duration of the G1-phase of GCPs in MCMV-infected animals as compared to GCPs in control uninfected animals. In contrast to reduced protein levels of cyclin E and Cdk2, we did not detect changes in the cyclin D1 or p-cyclin D1 levels in GCPs from infected mice while levels of Cdk4 and Cdk6 were increased and decreased, respectively, relative to control samples. Because the activities of Cdk4 and Cdk6 are redundant with regard to Rb phosphorylation, it is possible that the changes in the steady state levels of the Cdk catalytic subunits could have a negligible impact on its modification. Our results demonstrating unaltered levels of Rb phosphorylated on Ser795 or the presence of Rb phosphorylated Ser780 or Ser807/811, though significantly reduced in GCPs of MCMV-infected mice compared to non-infected control mice, argued that the cyclin D-Cdk4/6 complexes maintained kinase activity in GCPs of infected mice. These finding suggests that Rb phosphorylation is altered but still present in GCPs of MCMV-infected mice as compared to uninfected mice and further supports our findings that progression through G1-phase is delayed but not blocked in GCPs during MCMV infection.

Overexpression of cyclin D1 and cyclin E1 in the mouse cortical precursors resulted in a 25% reduction in the duration of G1-phase of the cell cycle, promoted cell cycle reentry, and inhibited neuronal differentiation (Pilaz et al., 2009). In contrast, lengthening of G1-phase, can cause proliferating cells to prematurely exit cell cycle, lead to neurogenesis, and subsequently result in the reduction of the overall population size of NPCs of the developing cerebral cortex (Calegari et al., 2005; Glickstein et al., 2009; Lanctot et al., 2017; Lange et al., 2009; Mitsuhashi et al., 2001; Suter et al., 2007). Our results are consistent with the findings in the NPCs, in that lengthening of the G1-phase was accompanied by increased cell cycle exit of GCPs of infected animals compared to non-infected control animals.

Collectively, our results argue that virus-induced inflammation and not direct virus cytopathic effects drive MCMV pathogenesis during postnatal cerebellar development. Importantly, our findings in MCMV-infected mice suggest that lengthening of the cell cycle of GCPs was due to prolonged G1- and S-phases and associated with premature cell cycle exit and delayed migration within the EGL. These findings together with further characterization of cellular responses to MCMV-induced inflammation could help identify specific targets that could contribute to the development of more efficacious therapeutic agents for the treatment of infants with congenital HCMV infection.

## Acknowledgements

This work was supported by a grant from NIH R01AI089956 (WJB)

## Author contribution

Conceptualization: Sung, C.Y.W., Jonjic, S., Britt, W.J. Data curation: Sung, C.Y.W., Li, M., Sanchez, V. Formal Analysis: Sung, C.Y.W. Funding acquisition: Britt, W.J. Investigation: Sung, C.Y.W., Li, M. Methodology: Sung, C.Y.W. Project administration: Sung, C.Y.W., Britt W.J. Resources: Jonjic, S., Britt, W.J. Software: Sung, C.Y.W. Supervision: Britt, W.J. Validation: Sung, C.Y.W. Visualization: Sung, C.Y.W. Writing – original draft: Sung, C.Y.W. Writing review & editing: Sung, C.Y.W., Britt, W.J.

---

**Supplemental Figure 1.**
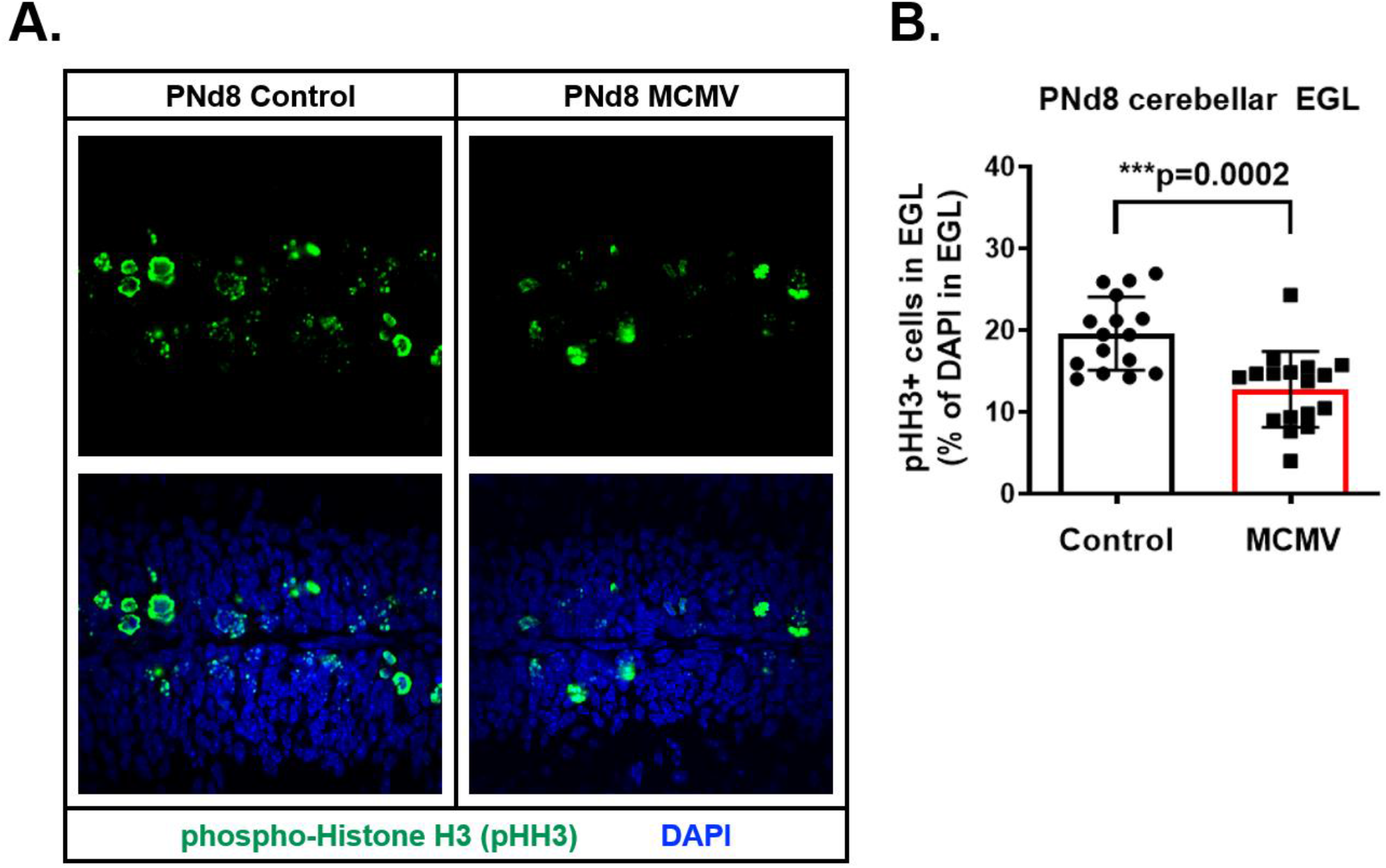
Fewer GCPs were in G2/M-phase of MCMV-infected cerebella. **(A)** Representative image of cerebella from non-infected control and MCMV-infected mice stained for phospho-histone H3 (pHH3) to quantify GCPs in G2/M-phase. **(B)** GCPs in G2/M-phase were quantified in percentage of pHH3^+^ GCPs to total number of nuclei in the EGL (pHH3^+^/total DAPI in the EGL). Data are shown as mean ± SD, n=3-4 mice/experimental group in three different regions in the cerebellum. P-values were calculated using two-tailed t test.

**Supplemental Figure 2.**
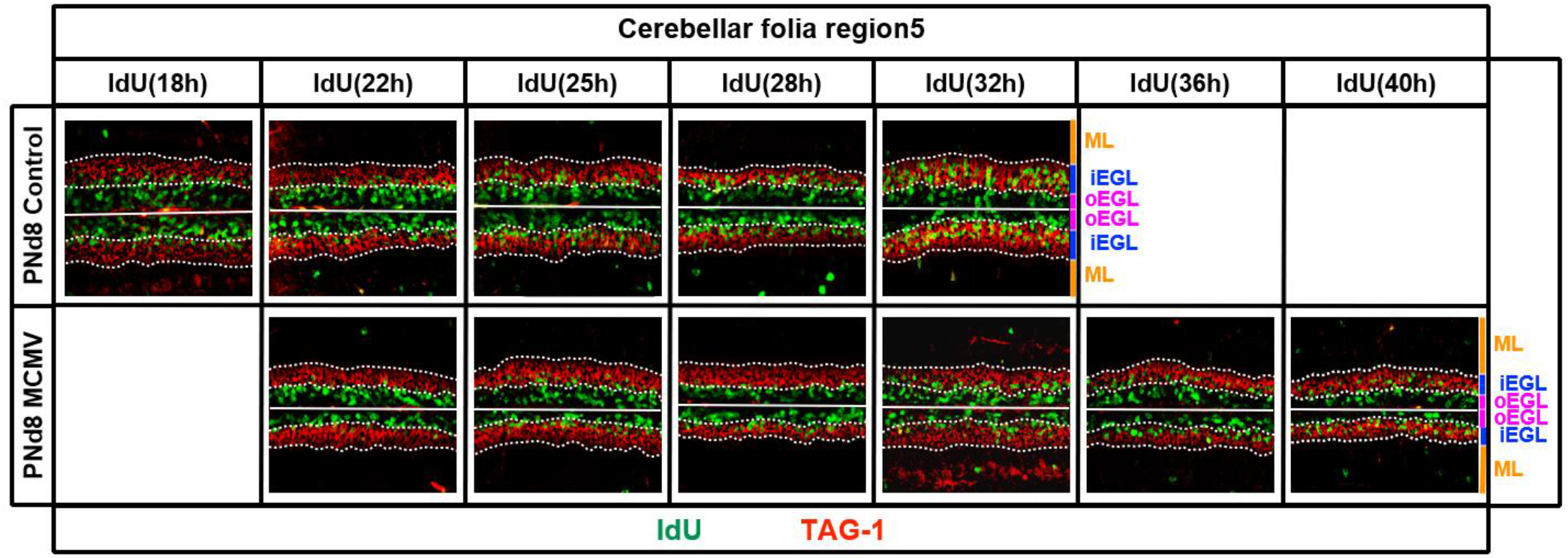
Migration of GCPs within the EGL is delayed in MCMV-infected cerebellum. Representative images of cerebella from non-infected and MCMV-infected mice stained for IdU (green) and TAG-1 (red) after treatment with IdU for 18, 22, 25, 28, 32, 36, and 40 hrs to track migration of GCPs. TAG-1 is a cell adhesion molecule highly expressed in axons of immature/mature GCs and specifically stains for the iEGL. Cerebellar folia are indicated by the white solid line and the iEGL is located between the two white dotted lines (TAG-1+ layer). Outer EGL (oEGL, magenta label); inner EGL (iEGL, blue label); molecular layer (ML, orange label). Data are representative images of n=4-6 mice/experimental group.

